# Functional binding dynamics relevant to the evolution of zoonotic spillovers in endemic and emergent *Betacoronavirus* strains

**DOI:** 10.1101/2020.09.11.293258

**Authors:** Patrick Rynkiewicz, Gregory A. Babbitt, Feng Cui, André O. Hudson, Miranda L. Lynch

## Abstract

Comparative functional analysis of the dynamic interactions between various *Betacoronavirus* mutant strains and broadly utilized target proteins such as ACE2 and CD26, is crucial for a more complete understanding of zoonotic spillovers of viruses that cause diseases such as COVID-19. Here, we employ machine learning to replicated sets of nanosecond scale GPU accelerated molecular dynamics simulations to statistically compare and classify atom motions of these target proteins in both the presence and absence of different endemic and emergent strains of the viral receptor binding domain (RBD) of the S spike glycoprotein. Machine learning was used to identify functional binding dynamics that are evolutionarily conserved from bat CoV-HKU4 to human endemic/emergent strains. Conserved dynamics regions of ACE2 involve both the N-terminal helices, as well as a region of more transient dynamics encompassing K353, Q325 and a novel motif AAQPFLL 386-92 that appears to coordinate their dynamic interactions with the viral RBD at N501. We also demonstrate that the functional evolution of *Betacoronavirus* zoonotic spillovers involving ACE2 interaction dynamics are likely pre-adapted from two precise and stable binding sites involving the viral bat progenitor strain’s interaction with CD26 at SAMLI 291-5 and SS 333-334. Our analyses further indicate that the human endemic strains hCoV-HKU1 and hCoV-OC43 have evolved more stable N-terminal helix interactions through enhancement of an interfacing loop region on the viral RBD, whereas the highly transmissible SARS-CoV-2 variants (B.1.1.7, B.1.351 and P.1) have evolved more stable viral binding via more focused interactions between the viral N501 and ACE2 K353 alone.

## INTRODUCTION

In the worldwide COVID-19 pandemic there has been an unprecedented wealth of high-throughput sequencing surveillance of the SARS-CoV-2 virus and near real-time sequence-based analysis of the evolution of local viral strains. Sequence-based analysis can be highly insightful for detailing the phylogenetic history of viral strains. However, it is often difficult to ascertain the functional aspects of this rapid viral evolution without expensive and time-consuming functional binding assays. These assays often need to follow highly specific biosafety level protocols when used with human pathogens or to use bioengineered protein constructs that are safer, but perhaps do not completely capture the biophysics of the complete pathogen. These functional assays include ELISA-based and solid phase assays for receptor binding specificity as well as microcalorimetry to quantify binding affinity. They can be aimed at identifying and quantifying the functional repercussions of single non-neutral variants from among many more neutral variants generated in the normal process of rapid viral evolution. They can be further supplemented by sequence-based phylogenetic tests of selection as well as experimental and computational structural-functional analyses. However, even when employed in totality, these methods are still often incapable of fully deciphering the precise molecular functions that underlie viral transmission. Ultimately, viral transmission is the consequence of soft-matter dynamics responding to weak chemical bonding interactions at many potential sites on both the viral and putative target proteins. The unprecedented pandemic of SARS-CoV-2 brings with it many questions regarding the functional evolution of both emergent and endemic *Betacoronavirus* strains in humans from present and past zoonotic spillover events. It has been recently demonstrated in vitro that ACE2 is the target of SARS-CoV-2 across a broad variety of mammalian taxa (Zhao et al. 2020). With the current surveillance of new variants like the highly transmissible and more lethal strains of SARS-CoV-2 (B.1.1.7, B.1.351, and P.1), the comparative biophysical analysis of potentially conserved dynamic mechanisms of viral protein binding *in silico* can offer a new perspective on the functional consequences of viral evolution during a pandemic (Rambaut et al. 2020; Faria et al. 2021; Tegally et al. 2021).

The SARS-CoV-2 virus belongs to the class of positive-strand RNA viruses (genus *Betacoronavirus*) and is structurally defined by a nucleocapsid interior surrounded by the spike, membrane, and envelope proteins (Haan et al. 2000; Fung and Liu 2019). Structural membrane and envelope proteins are known to play a key role in virion assembly and budding from infected cells, making them potential targets for antiviral treatments (Haan et al. 2000). One prominent protein candidate for comparative studies of the propensity for viral transmission is the spike or S glycoprotein, which is responsible for initial viral contact with the target host receptor and subsequent entry into host cells. The spike protein consists of two homotrimeric functional subunits: the S1 subunit consisting of the viral receptor binding domain (RBD) interacting directly with the host, and the S2 subunit which directly fuses the virion to the host cellular membrane (Hoffmann et al. 2018; Shang et al. 2020). The S1 and S2 subunits of coronaviruses are cleaved and subsequently activated by type-II transmembrane serine proteases and the subtilisin-like serine protease furin, the latter of which evidently primes SARS-CoV-2 for cellular entry but not SARS-CoV-1 (Follis et al. 2006; Shang et al. 2020). The boundary between the S1 and S2 subunits forms an exposed loop in human coronaviruses SARS-CoV-2, hCoV-OC43, hCoV-HKU1, and MERS-CoV that is not found in SARS-CoV-1 (Hoffmann et al. 2020) and suggests a complex functional evolutionary landscape for emergent SARS-type coronaviruses.

The spike protein of SARS-CoV-2 has been demonstrated to directly and specifically target the human angiotensin converting enzyme 2 (ACE2) receptor, which has a role in the renin–angiotensin system (RAS) critical for regulating cardiovascular and renal functions (Oudit et al. 2003; Ou et al. 2017; Ou et al. 2020). ACE2 is a glycoprotein consisting of 805 amino acid residues that is the primary target for docking and viral entry in many coronavirus infections, including those belonging to SARS-CoV and human alpha-coronavirus NL63 (Hofmann et al. 2005). ACE2 is classified as a zinc metallopeptidase (carboxypeptidase) involved in cleaving a single amino acid residue from angiotensin I (AngI) to generate Ang1-9 (Kuba et al. 2005). In addition, ACE2 also cleaves a single amino acid residue from angiotensin II (Ang II) to generate Ang1-7 (Kuba et al. 2005). ACE2 receptors reside on the surfaces of multiple epithelial cells, with tissue distribution predominantly in the lung and gut linings, but are also found in heart, kidney, liver and blood vessel tissues. Given the dominant role ACE2 plays in SARS-CoV-2 viral entry, understanding how sequence and structural variations affect the molecular dynamics of ACE2/RBD interactions can provide information to guide the rationale design and development of therapeutics targeting this interaction. But it is important to note that SARS-CoV-2 also targets other human protein receptors including several neurolipins (Cantuti-Castelvetri et al. 2020; Daly et al. 2020) and perhaps the known targets of other viral relatives such as the MERS-CoV target protein CD26. Therefore, it is important to demonstrate at a molecular level, how such promiscuity in binding interaction is achieved by the viral RBD and how it may have contributed to zoonotic spillovers from other species to humans. The comparative dynamic modeling of present, past, and potential progenitor viral strain interactions to ACE2 and other known targets of past outbreaks like protein CD26, can also potentially illuminate the dynamics of molecular interactions that may facilitate zoonotic transmission to humans as well as the molecular evolution toward emergent and endemic human viral strains.

Here, we present a systematic approach to the comparative statistical analysis and machine learning classification of the dynamic motions of proteins generated from accelerated all-atom molecular dynamics simulations. This method is subsequently leveraged to characterize the functionally conserved binding signatures of viral strains on specific target proteins at single amino acid site resolution. We employ DROIDS 4.2/maxDemon 3.0, a machine learning assisted software tool designed by our lab to statistically analyze/visualize comparative protein dynamics and then subsequently identify (i.e. classify) protein regions with functionally conserved dynamics that can be discriminated by the machine learning from more random motions caused by thermal noise (Babbitt et al. 2018; Babbitt, Fokoue, et al. 2020). The software also now uses information theoretics to quantify the relative impact of variants on regions of conserved dynamics and to identify amino acid site coordination (i.e. allostery) related to functionally conserved dynamic states. We subsequently analyze molecular dynamics in models of different *Betacoronavirus* strain S spike glycoprotein interactions with their primary human target protein ACE2 as well as a likely progenitor target in bats, CD26. These strains include the endemic human strains of *Betacoronaviru*s (hCoV-OC43 and hCoV-HKU1), emergent strains responsible for recent SARS-like outbreaks (i.e. SARS-CoV-1, SARS-CoV-2 and MERS-CoV), as well as models of hypothetical interactions with the bat progenitor strain bat CoV-HKU4. We confirm two regions of binding interaction on ACE2 that appear pre-adapted from two precise and stable binding interactions of the bat progenitor strain with CD26 and are functionally conserved across varying evolutionary distances of the human evolving strains. We confirm important key sites of binding interaction at the ACE2 N-terminal helices, K353 and Q325 identified previously by Yan et al. (Yan et al. 2020) and Li et al. (F. Li et al. 2005; W. Li et al. 2005) In addition, we identify a potentially new ACE2 interaction site with the viral N501 that exhibits coordinated dynamic interplay along with K353 and Q325. This indicates that the original outbreak strain of SARS-CoV-2 has quite a significant degree of binding promiscuity in its interactions with ACE2 after its zoonotic spillover into the human population. Importantly, we demonstrate that the new variants (B.1.1.7, B.1.351 and P.1) have lost this more promiscuous binding in favor of a much more static and stable interaction with K353 alone. Lastly, we demonstrate that while all *Betacoronavirus* spillovers both past and present seem to rely upon conserved ACE2 binding in the same regions, the functional evolution of the molecular binding mechanisms in the current emergent SARS-CoV-2 outbreak is progressing quite differently within these conserved sites when compared to the evolution of past human endemic strains (Lubin et al. 2020).

## MATERIALS AND METHODS

### Molecular dynamic simulation – some background

A powerful tool for *in silico* analysis of protein-protein interactions is all atom accelerated molecular dynamics (aMD) simulation, which attempts to accurately sample the physical trajectories of molecular structures over time via traditional Newtonian mechanics played out over a modified potential energy landscape that encourages conformational transition states (Pierce et al. 2012). Molecular dynamics (MD) simulations are increasingly broadening these methods of inquiry by allowing further analysis of the rapid time scale dynamics of protein-protein interactions. Protein structures determined via x-ray crystallography have previously been used to provide insights into viral evolution by comparing angstrom scale distances and conformational similarities. MD simulations have already been employed to map the hydrogen bonding topology of the SARS-CoV-2 RBD/ACE2 interaction The goal was to explore the merits of drug repurposing, to characterize mutations, and to determine the stability of binding interactions between the SARS-CoV-2 RBD and several contemporary antiviral drugs (Ahamad et al. 2020; Ali and Vijayan 2020; Al-Khafaji et al. 2020; Mittal et al. 2020; Muralidharan et al. 2020; Wang et al. 2020; Yu et al. 2020). While conventional MD and aMD simulations can provide unprecedented resolution on protein motion over time, interpretations based upon comparisons of single MD trajectories are statistically problematic and difficult to compare. This makes the application of MD to comparative questions regarding functional binding or impacts of mutation quite challenging. Given recent increases in GPU processor power, the statistical comparison of large replicates of aMD simulations is now an available solution to this problem. Because MD trajectories of complex structures like proteins are inherently complex and chaotic, even if they are adequately resampled, they do not often fit traditional model assumptions of many parametric statistical tests. Therefore, reliable comparisons of molecular dynamics need to further be based upon statistical methods that can capture the multimodal distribution space of protein motion and assign statistical significance for divergence in dynamics without over-reliance on the assumptions of normality in the distribution of molecular dynamics over time.

To this end, our lab has developed DROIDS 4.2, a method and software tool designed to address these challenges in the statistical analysis and visualization of comparative protein dynamics (Babbitt et al. 2018; Babbitt, Fokoue, et al. 2020). DROIDS software includes maxDemon 3.0, a machine learning algorithm/pipeline for detecting functionally conserved dynamics in new/novel MD simulations after being trained upon the functional dynamic states defined in DROIDS. The package now also includes information theoretics designed to compare the relative impacts of genetic variation upon functionally conserved dynamics, as well as identify coordination between sites that are involving the conserved dynamics. This method has been successfully applied to the evolution of activation loop dynamics relevant to cancer drug hyperactivation in the B-raf kinase protein (Babbitt, Lynch, et al. 2020). Here, we have subsequently applied these methods to analyze hybrid models of different strains of *Betacoronavirus* S spike glycoprotein interactions with their primary human target protein ACE2 as well as a likely bat progenitor strain target CD26.

### PDB structure and hybrid model preparation

Structures of the two main human SARS variants of beta-coronavirus spike glycoprotein receptor binding domain (RBD) bound to human ACE2 receptor protein were obtained from the Protein Data Bank (PDB). These were SARS-CoV-1 (PDB: 6acg) (Song et al. 2018) and SARS-CoV-2 (PDB: 6m17) (Yan et al. 2020). Three additional hybrid models of viral RBD interaction with ACE2 consisting of human *Betacoronavirus* variants MERS-CoV (PDB: 5×5c, and PDB 4kr0) (Yuan et al. 2017), hCoV-OC43 (PDB: 6ohw) (Tortorici et al. 2019), and hCoV-HKU1 (PDB: 5gnb) (Ou et al. 2017) bound to ACE2 were generated by creating an initial structural alignment to the SARS-CoV-2 RBD/ACE2 model (PDB: 6m17) using UCSF Chimera’s MatchMaker superposition tool (Pettersen et al. 2004) followed by deletion of the SARS-CoV-2 RBD in the model and any structures outside of the molecular dynamics modeling space defined in Fig 1A, leaving only the viral RBD and ACE2 receptor domain. These hybrid models of ACE2 interaction included viral receptor binding domains from MERS-CoV (PDB:5×5c and 4kr0) (Yuan et al. 2017), HCoV-OC43 (PDB: 6ohw) (Tortorici et al. 2019), and HCoV-HKU1 (PBD: 5gnb) (Ou et al. 2017) bound to the ACE2 protein from the SARS-CoV-1 model. Unbound forms of the ACE2 structure were obtained by deleting the viral structure in PDB: 6m17 and performing energy minimization for 2000 steps and 10ns of equilibration of molecular dynamics simulation in Amber18 (Case et al. 2005; Götz et al. 2012; Pierce et al. 2012; Salomon-Ferrer et al. 2013) prior to setting up production runs for the sampling regime (described below). Loop modeling and refinement were conducted where needed using Modeller in UCSF Chimera (Sali and Blundell 1993; Fiser et al. 2000). A hybrid model of the bat HKU4 beta coronavirus interaction with ACE2 was similarly constructed using PDB: 4qzv (Wang et al. 2014), batCoV-HKU4 bound to the MERS target human CD26, as a reference for structural alignment to PDB: 6m17. Our hybrid models of the viral RBD consisted of the largely un-glycosylated region (represented in PDB: 6m17 from site 335-518). Only one glycan was removed using swapaa in UCSF Chimera at ASN 343 on the viral RBD located on the opposite side of the RBD-ACE2 interface (PDB: 6m17). Five glycans on the ACE2 receptor domain were also removed, none occurring near the binding interface (ASN 53, ASN 90, ASN 103, ASN 322, ASN 546). Single mutation models were also similarly constructed to examine the effects of single mutations of potentially large effect. These models are summarized in Table 1 and are available in a separate supplemental download file titled PDBmodels.zip.

**Figure 1.**
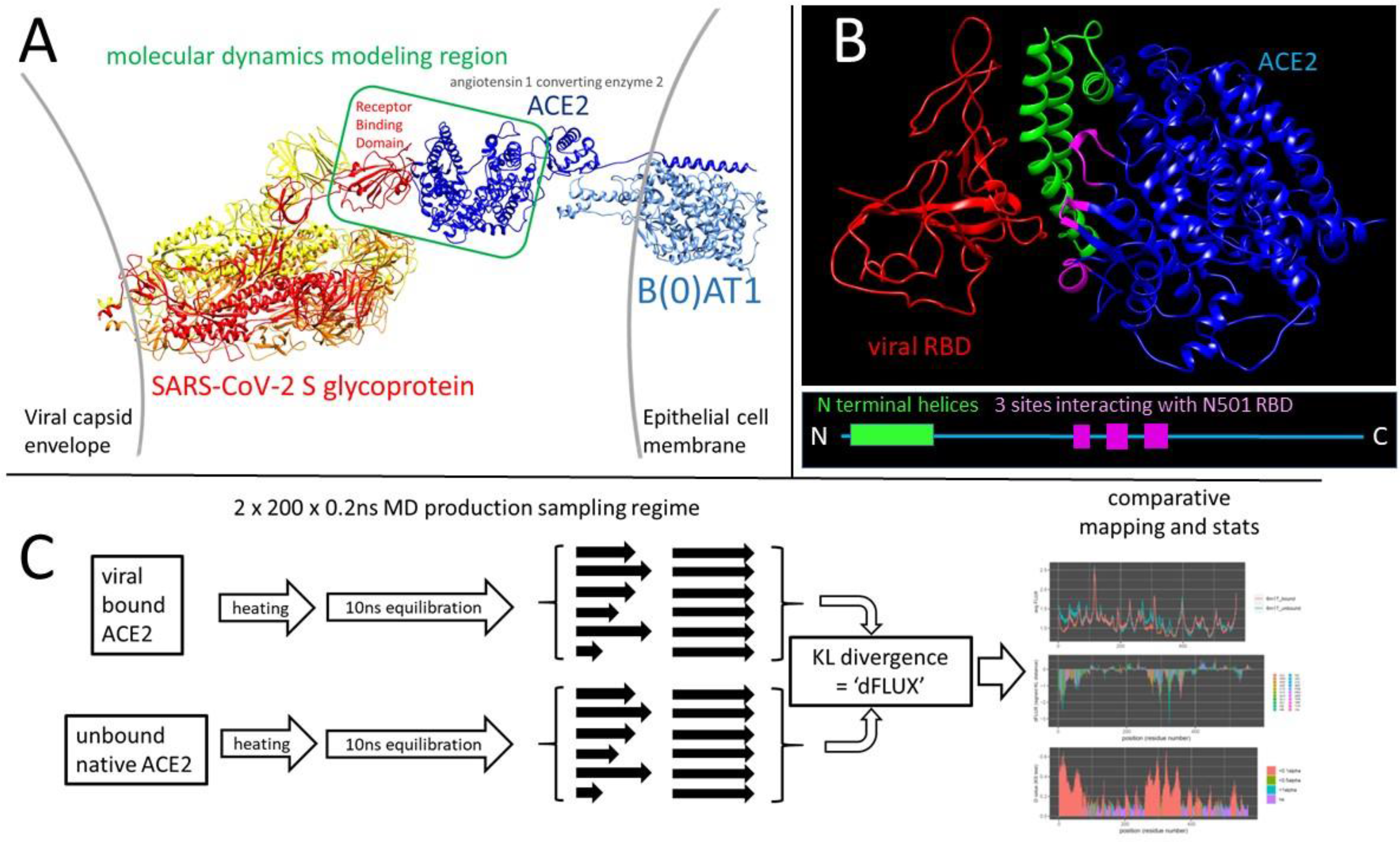
An overview of the molecular system and workflow to generate the comparative protein dynamics data sets. (A) The open conformation SARS-CoV-2 spike glycoprotein (PDB: 6vyb) interaction with its human target cell receptor, angiotensin 1 converting enzyme 2 (ACE2 PDB: 6m17). The green box in the center shows our modeling region that bounds the subsequent molecular dynamics (MD) simulations in the study. (B) The modeling region of the viral receptor binding domain (RBD) in red and ACE2 target protein in blue is shown along with the most relevant features of the binding interface; the N-terminal helices of ACE2 shown in green and the three main sites of the RBD N501 site interactions shown in magenta. (C) An overview of the workflow for the MD simulation pipeline for binding signature analysis used throughout this study. The difference in atom fluctuation or ‘dFLUX’ upon ACE2 binding by the virus is defined as the signed/symmetric KL divergence in the distribution of viral bound vs. unbound ACE2 atom fluctuations collected over 200 production runs and averaged to each amino acid site on the protein backbone (i.e. N, C, Cα, and O atoms).

**Table 1.**
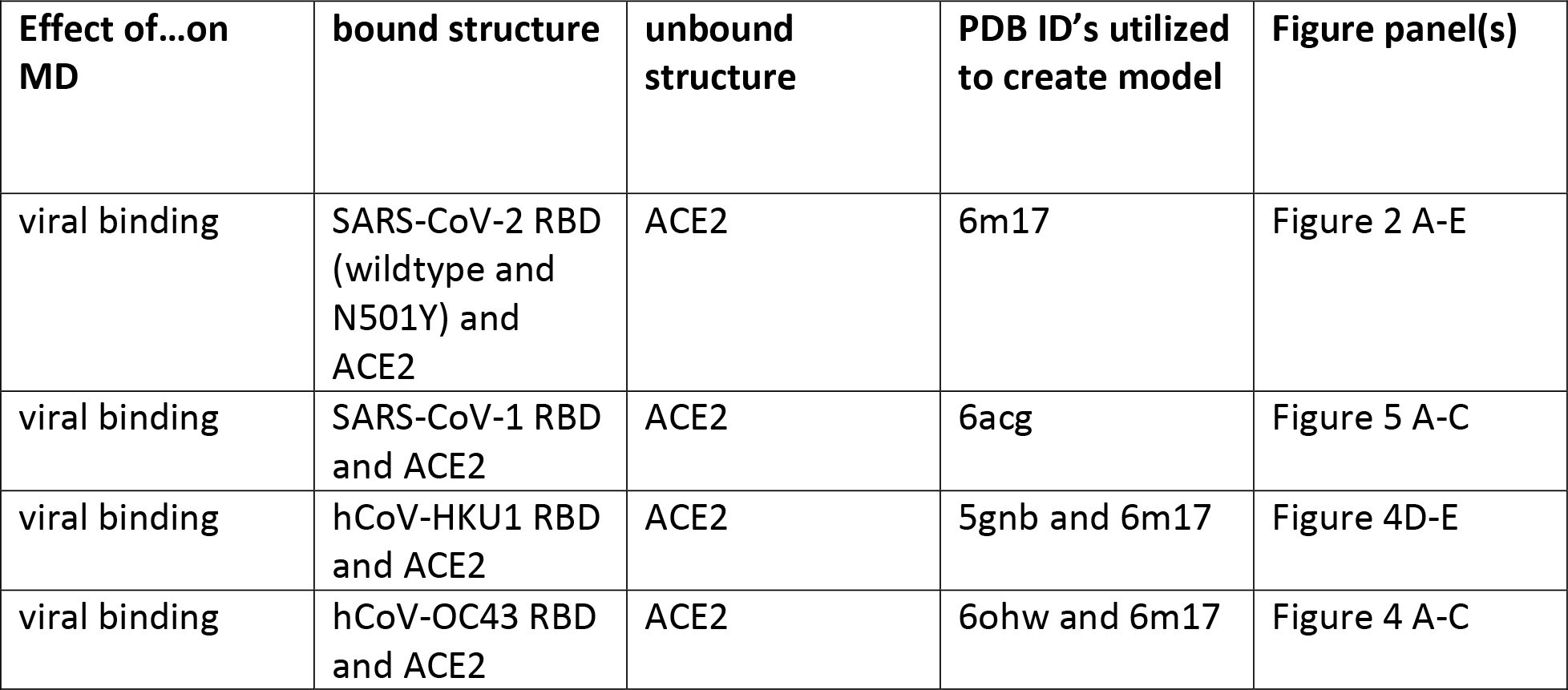
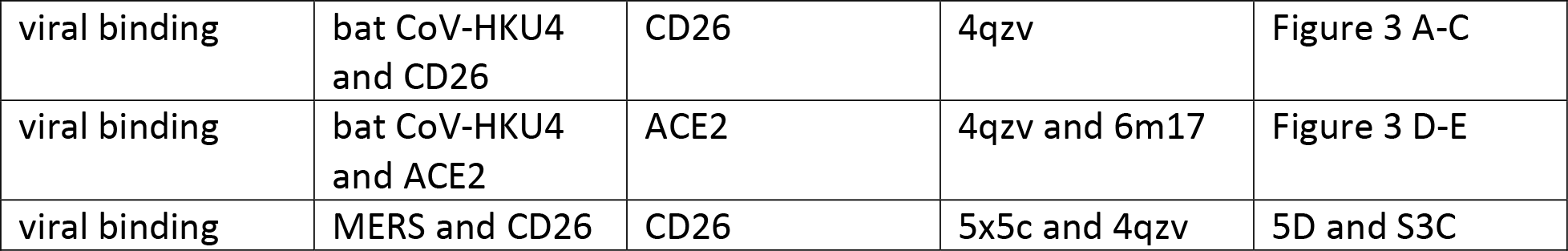
Hybrid models comparatively analyzed in this study.

### Mutant model construction and rationale

To generate *in silico* site-directed mutagenesis computations, we created mutant models using the swapaa function in UCSF Chimera 1.13 by first swapping amino acids using optimal configurations in the Dunbrack rotamer library, then using 2000 steepest gradient descent steps of energy minimization to relax the regions around the amino acid replacement. Mutant models were chosen to test the predicted sites functional role mitigating local binding between the viral RBD and ACE2 by comparing amino acid replacements with either very similar or very different sidechain properties. This is summarized in Table 2. The SARS-CoV-2 N501Y mutant model was constructed using PDB:6m17. This model of the viral RBD only extended from site 335-518 and therefore only included this single mutation (N501Y) from among the seven other amino acid replacements and two deletions found to occur on the B.1.1.7 lineage (The total set of mutations defining this variant are del 69-70, del 144, N501Y, A570D, D614G, P681H, T716I and S982A). Because of its proximity to the binding interface, the N501Y mutation is believed to be one of the most functional. In the B.1.351 and P.1 lineages, we were able to model three mutations in the viral RBD. These were N501Y, E484K and B417T/N.

**Table 2.**
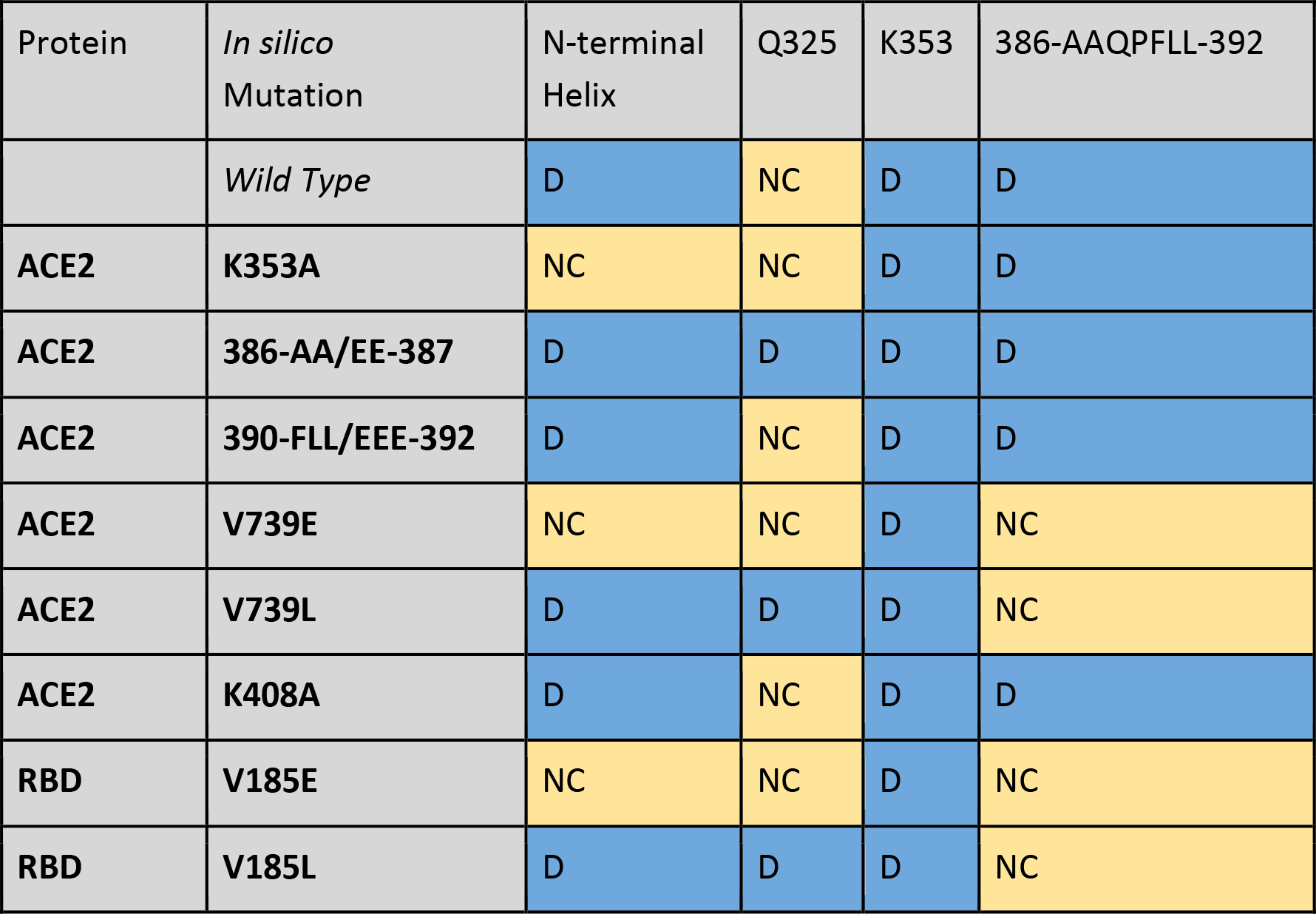
In silico targeted mutagenesis studies of the SARS-CoV-2 receptor-spike interaction with various amino acid mutations. Mutations were conducted with the *swapaa* command in UCSF Chimera, and bound structures were minimized with 2000 steps of steepest descent. D (blue) signifies dampened molecular motion, NC (yellow) indicates no significant change in molecular motion as a result of binding, and A (orange) indicates an amplification in molecular motion after RBD binding. Significance criteria was determined as having <= −1 KL divergence or less (dampened motion during binding), or >=1 KL divergence or more (amplified motion during binding). N-terminal helix and AAQPFLL motif categorization was determined as dampened or amplified when most of their constituent amino acids had a KL divergence of <= −1 or >= 1, respectively.

### Molecular dynamics simulation protocols

To facilitate the proper statistical confidence of the differences in rapid vibrational states in our model comparisons, many individual graphic processing unit (GPU) accelerated molecular dynamics (MD) simulations were produced in order to build large statistical replicate sets of MD trajectories. All accelerated MD simulations were prepared with explicit solvation and conducted using the particle mesh Ewald method employed by pmemd.cuda in Amber18 (Ewald 1921; Darden et al. 1993; Case et al. 2005; Pierce et al. 2012; Salomon-Ferrer et al. 2013) via the DROIDS v3.0 interface (Detecting Relative Outlier Impacts in Dynamic Simulation) (Babbitt et al. 2018; Babbitt, Fokoue, et al. 2020). Simulations were run on a Linux Mint 19 operating system mounting two Nvidia Titan Xp graphics processors. Explicitly solvated protein systems were prepared using tLeap (Ambertools18) using the ff14SB protein force field (Maier et al. 2015) and protonated at all available sites using pdb4amber in Ambertools18. Solvation was generated using the Tip3P water model (Mao and Zhang 2012) in a 12nm octahedral water box and subsequent charge neutralization with Na+ and Cl-ions. All comparative molecular dynamics binding analyses utilized the following protocol. After energy minimization, heating to 300K, and 10ns equilibration, an ensemble of 200 MD production runs each lasting 0.2 nanoseconds was created for both viral bound and unbound ACE2 receptor (Fig 1B). Two examples of the 10ns RMSF equilibration are also shown in Fig 1C. Each MD production run was preceded by a single random length short spacing run selected from a range of 0 to 0.1ns to mitigate the effect of chaos (i.e. sensitivity to initial conditions) in the driving ensemble differences in the MD production runs. All MD was conducted using an Andersen thermostat under a constant pressure of 1 atmosphere (Andersen 1980). Root mean square atom fluctuations (rmsf) and atom correlations were calculated using the atomicfluct and atomiccorr functions in CPPTRAJ (Roe and Cheatham 2013).

### Comparative dynamics of bound/unbound and mutant/wild type functional states

Computational modeling of bound and unbound protein structure dynamics can identify both proximal and distal differences in molecular motion as a result of the binding interaction. In the interest of studying functional dynamics, we focus on regions with highly dampened molecular motion at the interface between the viral RBDs and ACE2. Given the chaotic nature of individual molecular dynamics simulations, ensembles on the order of hundreds of samples are required for comparing functional states and deriving statistically significant differences between them. Furthermore, binding interactions can potentially be disrupted or prevented entirely by dynamic thermal noise that exists inherently in the system. DROIDS software employs a relative entropy (or Kullback-Leibler divergence) calculation on atom fluctuation distributions derived from large numbers of replicates of MD trajectories to quantify the local differences in both the magnitude and shape of the distribution of atom fluctuations in two different functional states of a protein. In this application, we compare viral bound versus unbound dynamics of target proteins to ascertain a computational signature of the local degree of dampened target protein motions during viral binding (Fig 2). DROIDS software also employs a non-parametric two sample Kolmogorov-Smirnov statistical test of significance, subsequently corrected for false discovery rates, to identify where these dampened dynamics upon viral binding are truly and locally significant. In our study, the dampened atom fluctuations of target proteins ACE2 and CD26 upon binding are indicative of molecular recognition, in the sense that weak bonding interactions overcome baseline thermal motion and lead to a persistent functional state.

**Figure 2.**
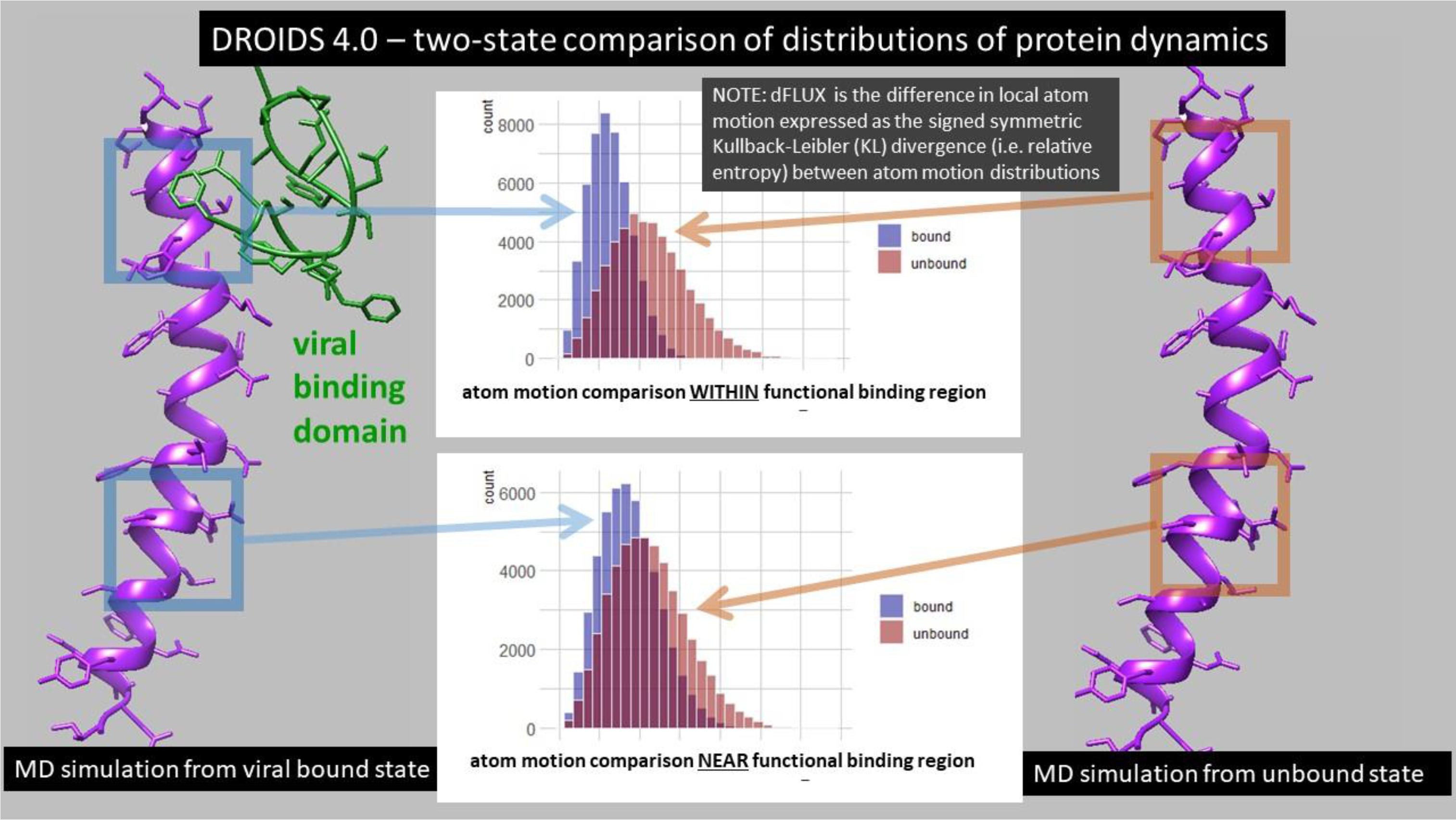
An overview of the comparative protein dynamics analysis with DROIDS 4.0. Two comparative functional protein states are defined and subjected to repeated sampling of molecular dynamics (MD) simulations. The differences in local atom motions (dFLUX) are defined by Kullback-Leibler (KL) divergences in protein backbone specific distributions in root mean square atom fluctuations (rmsf) collected over the MD sampling of each functional state. In this study, we compare the target protein human ACE2 in both its viral bound (blue-left) and its native unbound states (red-right). The reduced energy of motion at or near functional binding sites of the various viral strains will negatively shift the velocity of atoms creating a corresponding negative KL divergence in statistical distribution of rmsf (i.e. negative dFLUX).

The molecular dynamics of the viral bound and unbound models of ACE2/ were generated and statistically compared using the DROIDS v4.2 comparative protein dynamics software interface for Amber18 (Babbitt et al. 2018; Babbitt, Fokoue, et al. 2020). The symmetric Kullback-Leibler (KL) divergence or relative entropy (Kullback and Leibler 1951) between the distributions of atom fluctuation (i.e. root mean square fluctuation or *rmsf* taken from 0.01 ns time slices of total MD simulation time) on viral bound and unbound ACE2 were computed using DROIDS and color mapped to the protein backbone with individual amino acid resolution to the bound structures using a temperature scale (i.e. where red is hot or amplified fluctuation and blue is cold or dampened fluctuation). The *rmsf* value is thus

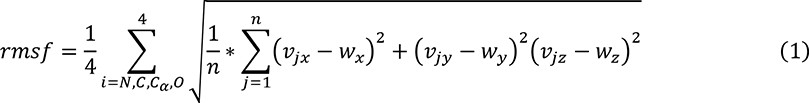

where v represents the set of XYZ atom coordinates for i backbone atoms (C, N, O, or Cα) for a given amino acid residue over j time points and w represents the average coordinate structure for each MD production run in a given ensemble. The Kullback Leibler divergence (i.e. relative entropy) or similarity between the root mean square fluctuation (rmsf) of two homologous atoms moving in two molecular dynamics simulations representing a functional binding interaction (i.e. where 0 = unbound state and 1 = viral bound state) can be described by

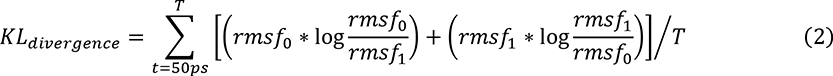

where rmsf represents the average root mean square deviation of a given atom over time. More specifically, the rmsf is a directionless root mean square fluctuation sampled over an ensemble of MD runs with similar time slice intervals. Because mutational events at the protein level are typically amino acid replacements, this calculation is more useful if applied to resolution of single amino acids rather than single atoms. Because only the 4 protein backbone atoms (N, Cα, C and O) are homologous between residues, the R groups or side chains are ignored in the calculation and equation 2 can be applied. Since the sidechain atoms always attach to this backbone, rmsf still indirectly samples the dynamic effect of amino acid sidechain replacement as they are still present in the simulation. The reference state of the protein is unbound while the query state is viral bound. Therefore, this pairwise comparison represents the functional impact of viral binding on the ACE2 protein’s normal unbound motion, where it is expected that viral contact would typically dampen the fluctuation of atoms at the sites of binding to some degree. This calculation in equation 2 is used to derive all the color maps and KL divergence (i.e. dFLUX) values presented in the Figures. Multiple test-corrected two sample Kolmogorov-Smirnov tests are used to determine the statistical significance of local site-wise differences in the *rmsf* distributions in viral bound and unbound ACE2 models. The test was applied independently to each amino acid site. The Benjamini-Hochberg method was used to all the adjust p-values for the false discovery rate generated from the multiple sites of a given protein structure.

### Machine learning to detect functionally conserved binding dynamics

We employed maxDemon 3.0 (Fig 3) for the machine learning detection of the functionally conserved dynamics of the binding interaction between SAR-CoV-2 and ACE2, the viral bound and unbound ACE2 structures modeled from PDB ID: 6m17 were each subjected to MD sampling consisting on 100 production runs each for 1.0 ns time, after 1ns heating and 10ns equilibration. Each production run was subdivided into short 50 frame time slices, from which subsequent atom fluctuation and atom correlations were computed using the atomicfluct and atomiccorr functions in cpptraj (Roe and Cheatham 2013) with a single residue mask applied only to the 4 atom types on protein backbone (N. Cα, C, O). The atom correlations for a given amino acid site were parsed only to include correlations to downstream sites at 1, 3, 5, and 9 positions distant. The pre-classifications of each time slice were simply determined from whether they originated from the viral bound or unbound simulation of ACE2. Thus, a feature vector for training the machine learning model on a given site (Fig 3A) was comprised of the pre-classifications (0=unbound,1=viral bound), the atom fluctuations at the given site (flux0) and the downstream correlations (corr1, corr3, corr5 and corr9). Two new single MD simulations were conducted (1ns heating, 10ns equilibration and 3ns production) on the zoonotic progenitor model bat CoV-HKU4 bound to human ACE2, and SARS-CoV-2 bound to ACE2. These two MD runs were subjected to time slicing and feature extraction as in the training set and then were used as validation runs for the machine learning. Because they represent the dynamic states of evolutionary orthologs, any local similarity of learning performance shared between them will represent functional binding dynamics, defined in the training set, that has persisted and been conserved over the time of evolutionary divergence of the virus. A multi-agent or ‘stacked’ model of shallow learners was individually applied to each amino acid site in the ortholog validation runs and learning performance profiles for each learning method were calculated as the average classification state over all the time slices (NOTE: a learning performance = 0.5 indicates no functional dynamics is detected at that position). An example of the learning performance profiles for all seven learners is shown for one of the validation runs in Fig 3B. The classification methods included K-nearest neighbors, naïve Bayes, linear and quadratic discriminant analysis, support vector machine with radial basis function kernel, random forest with 500 decision trees, and adaptive gradient boosting. All machine learning classifications are conducted in R using the class, MASS, kernlab, randomForest, and ada packages (Breiman 2001; Venables and Ripley 2010; Culp 2019; Karatzoglou 2019). Functionally conserved viral binding dynamics between the orthologs was captured using a canonical correlation analysis (Härdle and Simar 2007) applied to the 14 learning profiles (i.e. 7 for each ortholog in red and blue; Fig 3C) using a sliding window of 30 sites and an alpha level of significance of 0.01. An example of the local R-value and regions of significance (dark grey) are shown in Fig 3D. For more details, see our software publications (Babbitt et al. 2018; Babbitt, Fokoue, et al. 2020).

**Figure 3.**
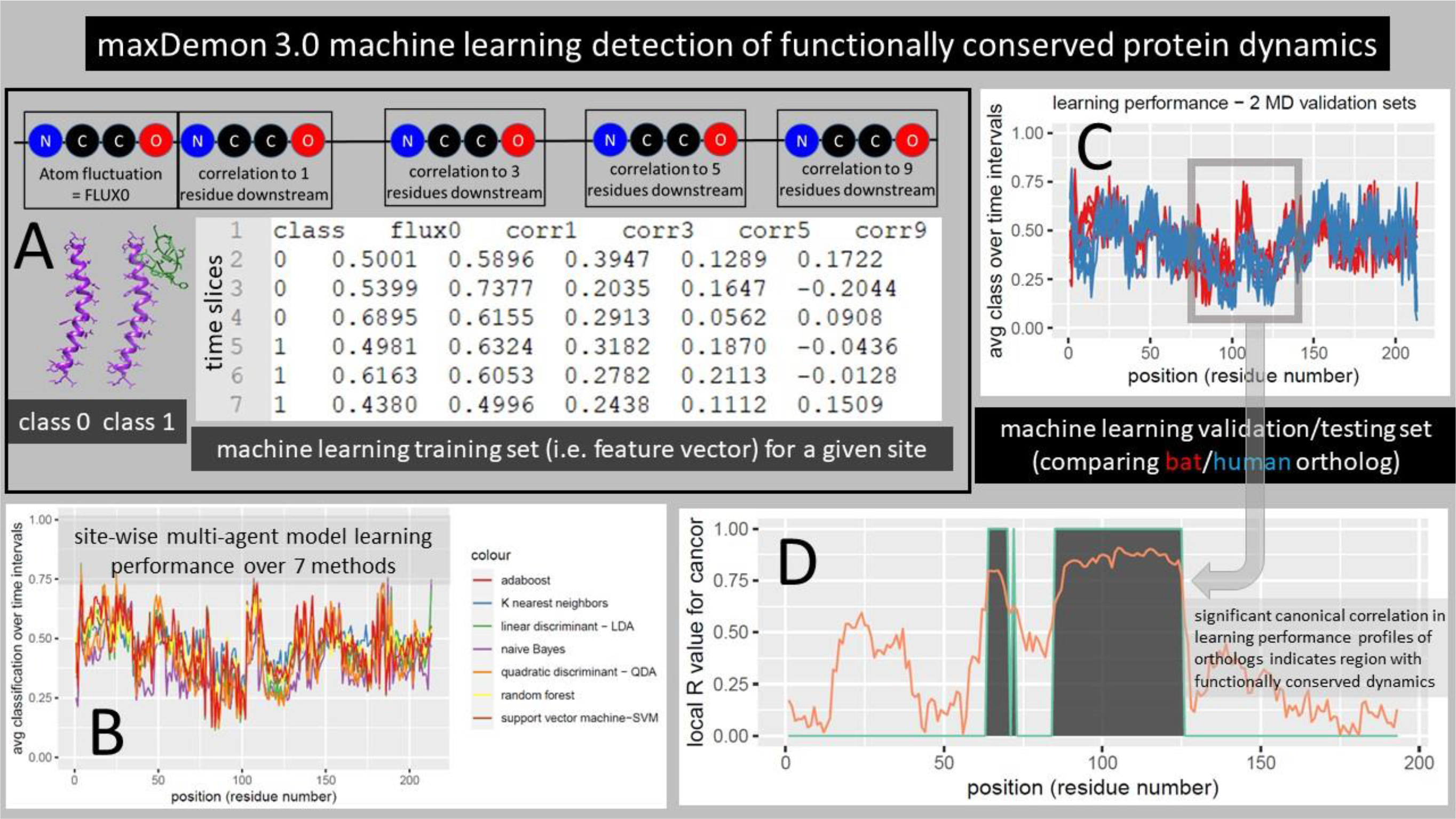
Detecting functionally conserved dynamics with the maxDemon 3.0 machine learning algorithm. (A) The MD simulation sets generated above are now the source of a pre-classified machine learning training set where class 0 and class 1 represent the binary functional states of the protein interaction (unbound vs bound). A feature vector is constructed comprising the atom fluctuations of any given amino acid (AA) site and its correlations to motions of downstream sites located 1, 3, 5, and 9 AA sites distant taken at every 50 frame interval or ‘time slice’ in the MD simulations. This rich feature vector thus captures both the local energy on the protein backbone, as well as potential sequence-based information that coordinates specific motions with nearby AA sites. (B) A multi-agent classifier comprised of up to seven learning methods is individually deployed on each site and a learning performance profile across all sites on the protein is generated as the time average classification of the site-specific dynamics defined by the feature vector. Thus, successful learning of functional dynamic features will deviate from an average classification of 0.5 on the profile. (C) This learning profile is then modeled for two evolutionary homologs (e.g. human ACE2 bound to the recent emergent strain SARS-CoV-2 and bound to bat CoV-HKU4, a potential zoonotic precursor) (D) A local canonical correlation of the learning performance profiles from both homologs is taken over 20 adjacent AA sites from a sliding window moved across the protein. Significant correlation of the homologous learning profiles identifies regions (shown in gray) where the functional binding dynamics defined by the pre-classification is functionally conserved from the time of divergence of the viral homologs.

### Information theoretics to detect variant impact and amino acid site coordination in conserved dynamics

After the functionally or evolutionary conserved dynamic regions are determined, the DROIDS/maxDemon software offers two new information theoretical calculations that can be applied to subsequent MD simulations conducted upon genetic or drug class/ligand binding variants of interest. The first metric is an information gain or relative entropy of correlation (REC) which can be used to compare the relative impacts of different variants on the functionally conserved dynamics defined in the section above (Fig 4A). This is a relative entropy calculation comparing the R values of the canonical correlations of each variant to that of the homologous functional validation described in the section above. REC calculations are returned locally in a sliding window as a line plot for each variants, and it is calculated globally over the whole protein backbone for each variant and returned as a bootstrapped bar plot with a one-way ANOVA test comparing all variants.

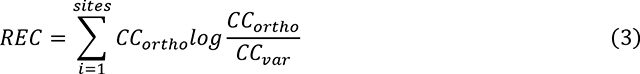

**Figure 4.**
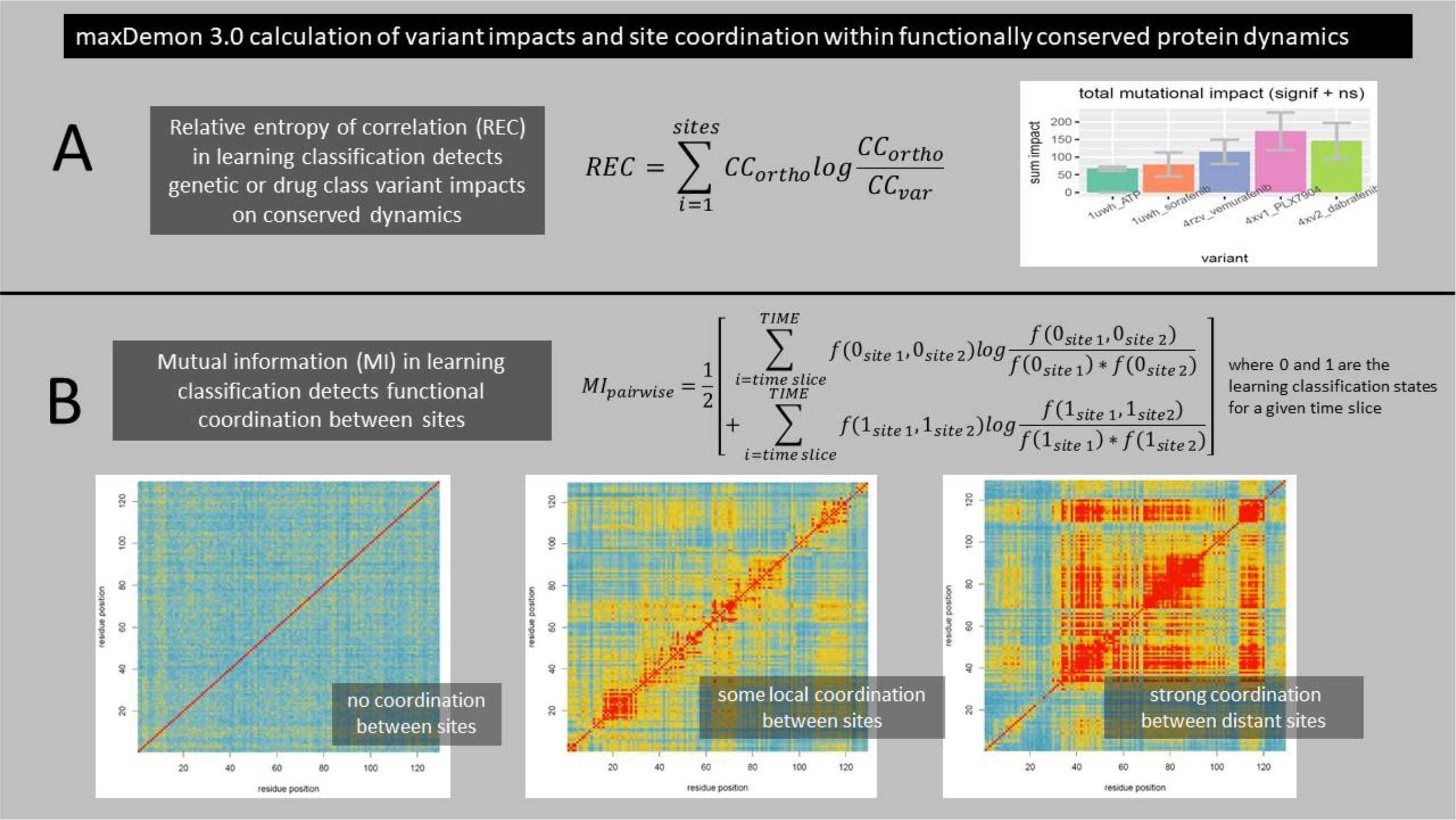
Information theoretics for measuring variant impacts and functional coordination between sites. Further information theoretics applied to the predicted binary classifications (0 or 1) of new MD runs can be used to compare relative impacts of different variants upon the signal of the conserved dynamics and to detect specific sites of functional coordination of conserved dynamics on the protein. (A) A relative entropy of the correlation (REC) of learning performance profiles can be used to measure both the local and total impact of genetic variation in the viral receptor binding domain on the functionally conserved binding dynamics of the bat/human orthologs. (B) Mutual information between the predicted binary states of learning classification (0 or 1) between any two amino acid sites, collected over the time of the simulations, can be used to detect functionally relevant coordination of the binding dynamics between the two sites. The mutual information is reported for all pairwise comparisons on the ACE2 protein as a heatmap with a scale blue/yellow/red indicating increasing coordination of functional binding dynamics.

A second information theoretic used is a mutual information (MI) for all pairwise site comparisons calculated using the learning classifications of all the time slices in a subsequently deployed MD simulation (Fig 4B). This calculation captures the frequency of the co-occurrence of learning classifications at any two given sites compared to the independent frequencies of the learning classifications at the two sites.

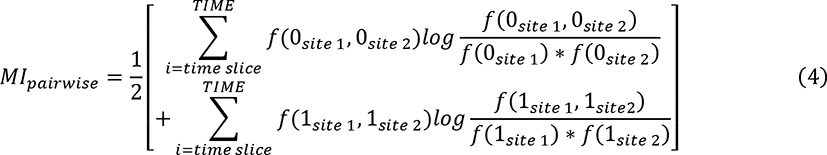

All site-wise comparisons are presented in a heatmap with blue/yellow/red scale that indicates the leve9l of co-occurrence of functional dynamic states. This provides a useful way of quantifying the amount of site coordination in functional dynamics which can be caused by allostery and/or cooperative binding interactions.

## RESULTS

### Characterization of the SARS-CoV-2 to ACE2 binding signatures in the original and N501Y mutant strain

The modeled region of binding interaction consisted of the extracellular domain region of human ACE2, and the viral RBD specified in Fig 1A. The main key sites of the ACE2 interface with the viral RBD are shown in Fig 1B, with the N-terminal helices labeled in green and the three main sites that can potentially interact with the viral RBD N501 site are shown in magenta. The DROIDS pipeline (Babbitt et al. 2018; Babbitt, Fokoue, et al. 2020) used to characterize the changes in ACE2 molecular dynamics due to viral binding is summarized Fig 1C. More details about this can be found in the methods section here as well as in our two software notes cited above. All the binding models developed and analyzed in this study are listed in Table 1. Comparisons of the molecular dynamics of SARS-CoV-2 RBD bound ACE2 to unbound ACE2 after long range equilibration revealed extensive dampening of atom fluctuations (shown color mapped in blue) at the ACE2 N-terminal helices as well as three downstream sites on ACE2 focused at Q325, K353, and a novel motif AAQPFLL 386-92 (Fig 5A-C). These color mapped structures show the local site-wise mathematical divergence protein backbone atom fluctuation (i.e. root mean square fluctuation averaged over N, C, Cα, and O) when comparing viral bound ACE2 dynamics to unbound ACE2 dynamics. They are presented for both the whole structural model (Fig 5A) and the viral binding interface with ACE2 (Fig 5B) along with sequence-based positional plotting of the signed symmetric Kullback-Leibler or KL divergence (Fig 5C). The site-wise positional profiles of average atom fluctuation in both the viral bound and unbound states (Fig S1), as well as the multiple test-corrected statistical significance of these KL divergences determined by a two-sample Kolmogorov-Smirnov or KS test is also given (Fig S2). NOTE: the false discovery rate for the p-value adjustment of each test was determined using the Benjamini-Hochberg method to adjust for the fact that a separate p-value is generated for each amino acid in the sequence. Therefore, the p-value given at each site is adjusted for the probability of false discovery that is governed by the length of the polypeptide.

**Figure 5.**
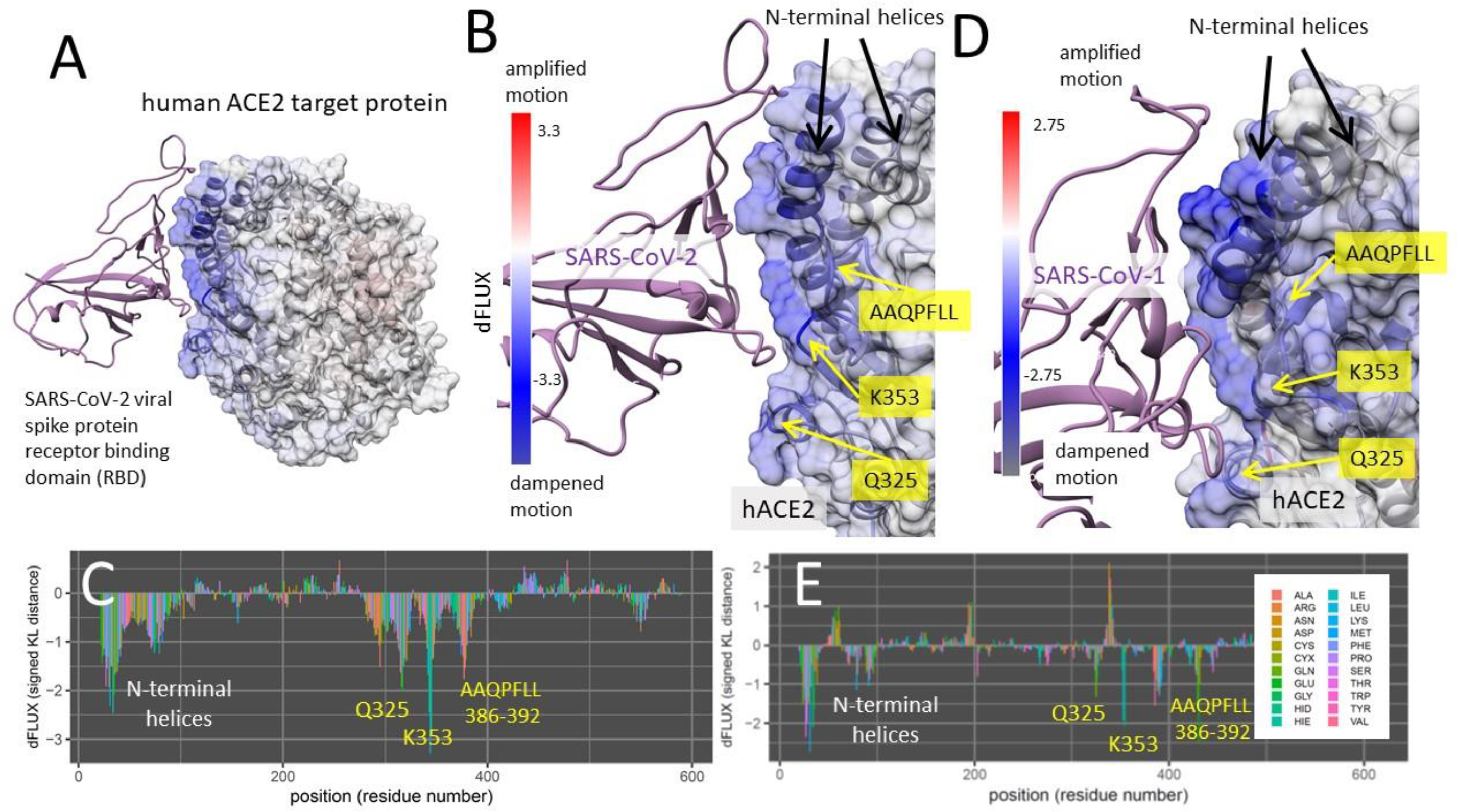
DROIDS binding signature of dampened atom fluctuations in human ACE2 receptor protein upon interaction with both SARS-CoV-2 spike glycoprotein (modeled from PDB: 6m17) and the upon interaction with the past human outbreak strain SARS-CoV-1 spike glycoprotein (modeled from PDB: 6acg). Shift in atom fluctuations were calculated as the signed symmetric Kullback-Leibler (KL) divergence (i.e. relative entropy or dFLUX) of distributions of the root mean square time deviations (rmsf) for residue averaged protein backbone atoms (i.e. N, Cα, C and O) for ACE2 in its spike bound (PDB: 6m17/6acg) versus unbound dynamic state (PDB: 6m17/6acg without the viral receptor binding domain or RBD). Here we show color mapping (A, B, D) and sequence positional plotting (C, E) of dampening of atom motion on viral RBD-protein target interface in blue for (A-C) the original strain of SARS-CoV-2 and (D-E) its predecessor, SARS-CoV-1. The sequence profile of the KL divergence between SARS-CoV-2 viral bound and unbound ACE2 produces strong negative peaks indicating key residue binding interactions with the N-terminal helices (white) and the N501 interactions at Q325, K353 and the AAQPLL 386-392 motif (yellow). The interactions at these sites are more moderated in the SARS-CoV-1 interaction with the ACE2 target protein.

Binding interaction models of past human outbreak strain SARS-CoV-1 in complex with ACE2 (Fig 5D-E) indicates that the binding signature of SARS-CoV-1 and SARS-CoV-2 are nearly identical with both having stable interaction with N-terminal helices of ACE2 and promiscuous interactions with K353 and the same novel sites Q325 and AAQPFLL 386-92. However, the SARS-CoV-1 interactions at these sites appear more moderated than in SARS-CoV-2. Again, the site-wise atom fluctuation profiles and KS tests of significance of the differences in these fluctuations are also given (Fig S1 C-F and S2 C-F).

### Characterization of the SARS-CoV-2 to ACE2 binding signatures in the N501Y variant strains B.1.1.7 and B.1.351

An identical characterization of the ACE2 binding signature created by the reportedly more highly transmissible SARS-CoV-2 B.1.1.7 (UK) variant indicated a similar yet stronger binding profile at the N-terminal helices, with more attraction to the C helix (Fig 6A-C). The binding to the ACE2 K353 by the viral RBD Y501 was also far more stable while the former promiscuous interactions with Q325 and AAQPFLL 386-92 were markedly reduced (Fig 6B-C, with point of the viral RBD mutation N501Y marked by the green star in Fig 3B). In the B.1.351 (South African) variant, we also find a more stable binding pattern overall that involves both the N-terminal helices and K353 site in ACE2 (Fig 6D-E). In this variant, we observed a retention of promiscuous binding of Y501 with the k353, Q325 and the AAQPFLL motif as well (Fig 6E). The three mutations prevalent in the RBD of the B.1.351 and P.1 variant are shown with a green star in Fig 6D. These are N501Y, E484K and K417N/T. Both variants display an extraction of the K353 side chain away from the body of ACE2 and towards the viral RBD Y501 (Fig 6B and 6D) that is generally maintained throughout the MD simulations.

**Figure 6.**
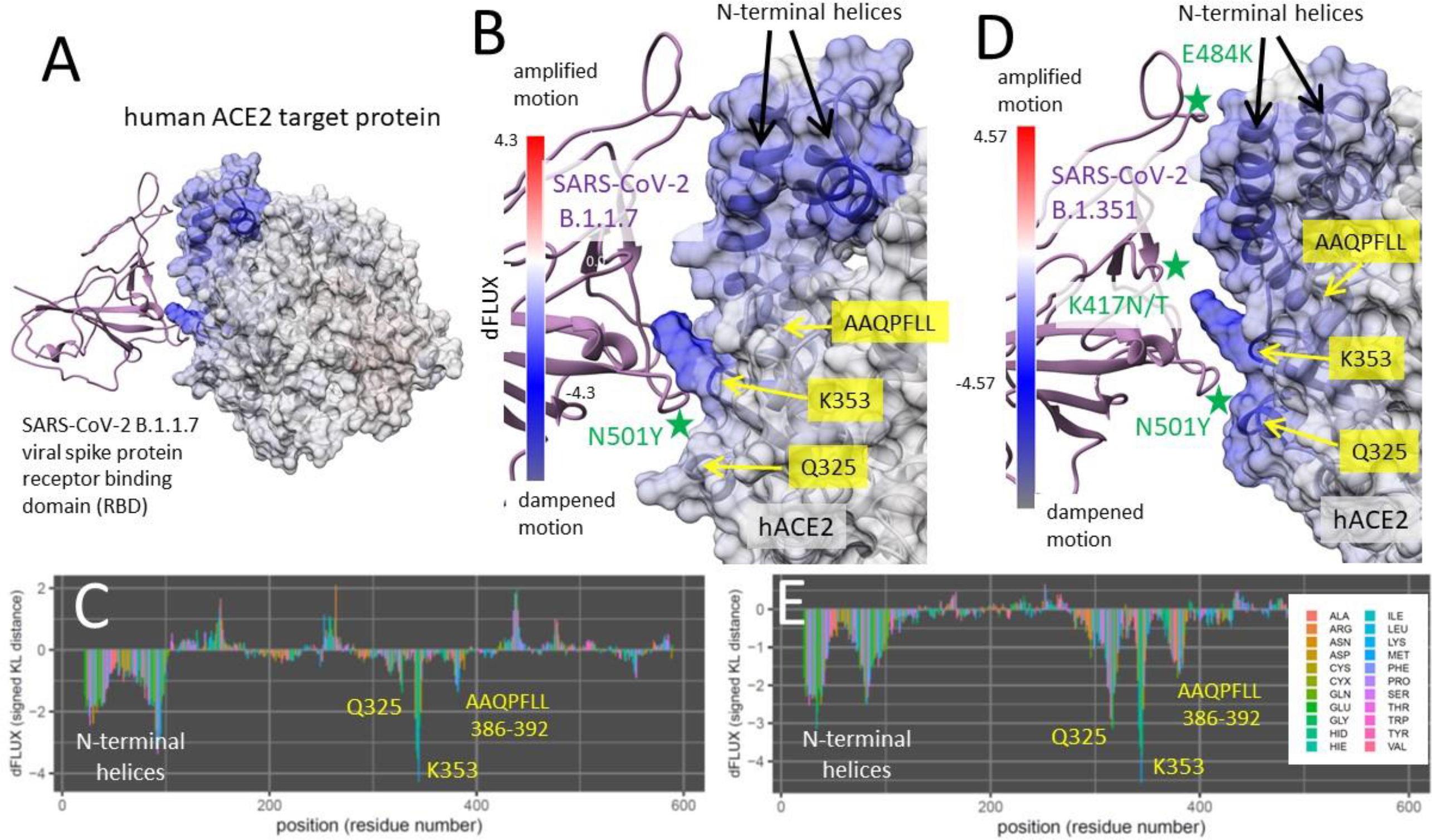
DROIDS binding signature of dampened atom fluctuations in human ACE2 receptor protein upon interaction with both the highly transmissible B.1.1.7 (UK) variant and the B.1.351 (South African and Brazilian) variant of SARS-CoV-2 spike glycoprotein (modeled from PDB: 6m17). Shift in atom fluctuations were calculated as the signed symmetric Kullback-Leibler (KL) divergence (i.e. relative entropy or dFLUX) of distributions of the root mean square time deviations (rmsf) for residue averaged protein backbone atoms (i.e. N, Cα, C and O) for ACE2 in its spike bound (PDB: 6m17) versus unbound dynamic state (PDB: 6m17 without the viral receptor binding domain or RBD). Here we show color mapping (A, B, D) and sequence positional plotting (C, E) of dampening of atom motion on viral RBD-protein target interface in blue for (A-C) the B.1.1.7 variant and (D-E) the B.1.351 variant of SARS-CoV-2. The sequence profile of the KL divergence between viral bound and unbound ACE2 produces negative peaks indicating key residue binding interactions with the N-terminal helices (white) and the N501 interactions at Q325, K353 and the AAQPLL 386-392 motif (yellow). The N501Y, E484K and K417N/T mutations are marked by the green stars (2B and 2D).

### Characterization of binding signatures in human endemic strains and past emergent outbreaks

Binding interaction models of human endemic strains hCoV-OC43 and hCoV-HKU1 in complex with ACE2 were also analyzed identically to previous models. The interaction with the most pathologically benign and longest co-evolved strain hCoV-OC43 (Fig 7A-C) shows highly enhanced dampened motion due to binding only at the N-terminal helices of ACE2, while the interaction with hCoV-HKU1 demonstrates this as well, but also adds moderate promiscuous interactions of N501 with K353 and Q325 (Fig 7D-E). The enhanced binding at the N-terminal helix is clearly associated with the evolution of extended loop structure of the viral RBD at this region of the interface (Fig 7A). The site-wise atom fluctuation profiles and KS tests of significance of the differences in these fluctuations are also given (Fig S1 C-D and S2 C-D). Binding interaction models of past human outbreak strain MERS-CoV in complex with both CD26 and ACE2 (Fig S4) indicate a two-touch interaction with CD26 sites SAMLI 291-5 and SS 333-4 (Fig S4 B-C). The MERS-CoV/ACE2 interaction indicates a strong interaction with the N-terminal helices and only a weak interaction with K353 (Fig S4 D-E). In the MERS-CoV MD simulations, we observe a similar maintenance of beta sheet over the SAMLI 291-5 region in CD26, however this is partially lost over the N-terminal helices when the MERS-CoV is presented to ACE2.

**Figure 7.**
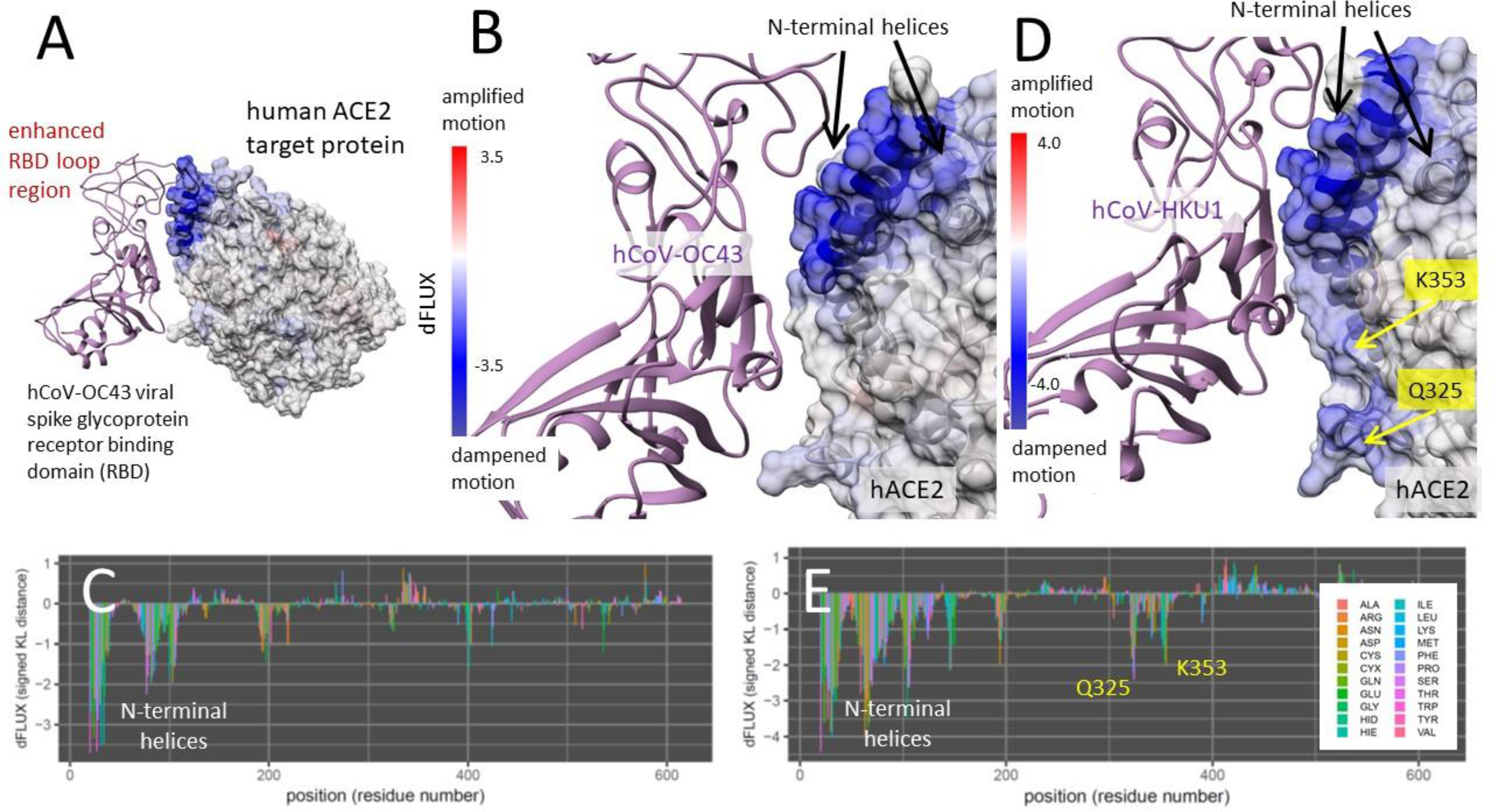
DROIDS binding signature of dampened atom fluctuations in human ACE2 receptor proteins upon interaction with two human commensal strains hCoV-OC43 and hCoV-HKU1 glycoprotein (modeled from PDB: 6ohw, 5gnb and 6m17). Here we show color mapping (A, B, D) and sequence positional plotting (C, E) of dampening of atom motion on the viral RBD-protein target interface in blue for (A-C) the targeting of ACE2 by the most benign strain hCoV-OC43 and (D-E) its less benign counterpart hCoV-HKU1. The sequence profile of the KL divergence between viral bound and unbound target proteins produces strong negative peak indicating key residue binding interactions (C, E) with the N-terminal helices on ACE2 and (E) moderate interactions with K353 and Q325 observed only in hCoV-HKU1.

We also examined the dampening of atom fluctuations in the viral receptor binding domains (RBD’s) of SARS-CoV-2 and the two human endemic strains (Fig S5). We find that in general, all of the viral RBD’s show stronger negative dFLUX than their ACE2 targets, indicative of their thermodynamic favoring of the target-bound state as one would expect in a viral system. Interestingly though, while we find that the SARS-CoV-2 RBD shows strong universal dampening across the whole RBD (Fig S5 A), the human strains demonstrate the strongest change in dynamics at the enhanced loop regions of the RBD (Fig S5 B-C), perhaps suggestive of the evolution of a disordered structure that becomes ordered only as it binds its target. Such a mechanism could obviously help the virus evade immune recognition, and we note that this has evolved in the same location in the endemic strains as the site of the E484K mutation that is currently linked to the ability of the SARS-CoV-2 variants that appear to have partially escaped vaccine. Also of interest, is a common small region of amplified atom fluctuation that occurs upon binding in all three RBD’s (labelled with orange question mark in Fig S5 A).

### Characterization of binding signatures of the bat progenitor strain HKU4 with its primary target CD26 and potential zoonotic spillover target ACE2

Binding interaction models of *Tylonycteris* bat progenitor strain batCoV-HKU4 RBD in complex with CD26 (its normal target) and ACE2 were also analyzed in the same manner as SARS-CoV-2 (Fig 8). The bat coronavirus HKU4 was strongly and significantly dampened molecular motion in two precise regions of CD26, the SAMLI 291-295 motif, and double serine motif at SS-333-4 (Fig 8A-C). However, upon interacting with ACE2, the bat coronavirus RBD only dampened the atom fluctuation at a broader region of the ACE2 N-terminal helices (Fig 8D-E) utilizing the same region that interacts with the CD26 SAMLI motif. Thus, the interaction with the ACE2 N-terminal helix appears pre-adapted by the evolution of the interaction with SAMLI on CD26. The site-wise atom fluctuation profiles and KS tests of significance of the differences in these fluctuations are also given (Fig S3). In both comparative MD simulations, we observe that the bat CoV-HKU4 viral RBD maintains more secondary structure in the form of anti-parallel beta sheet over the both the SAMLI 291-5 stie in CD26 and the N-terminal helices in ACE2.

**Figure 8.**
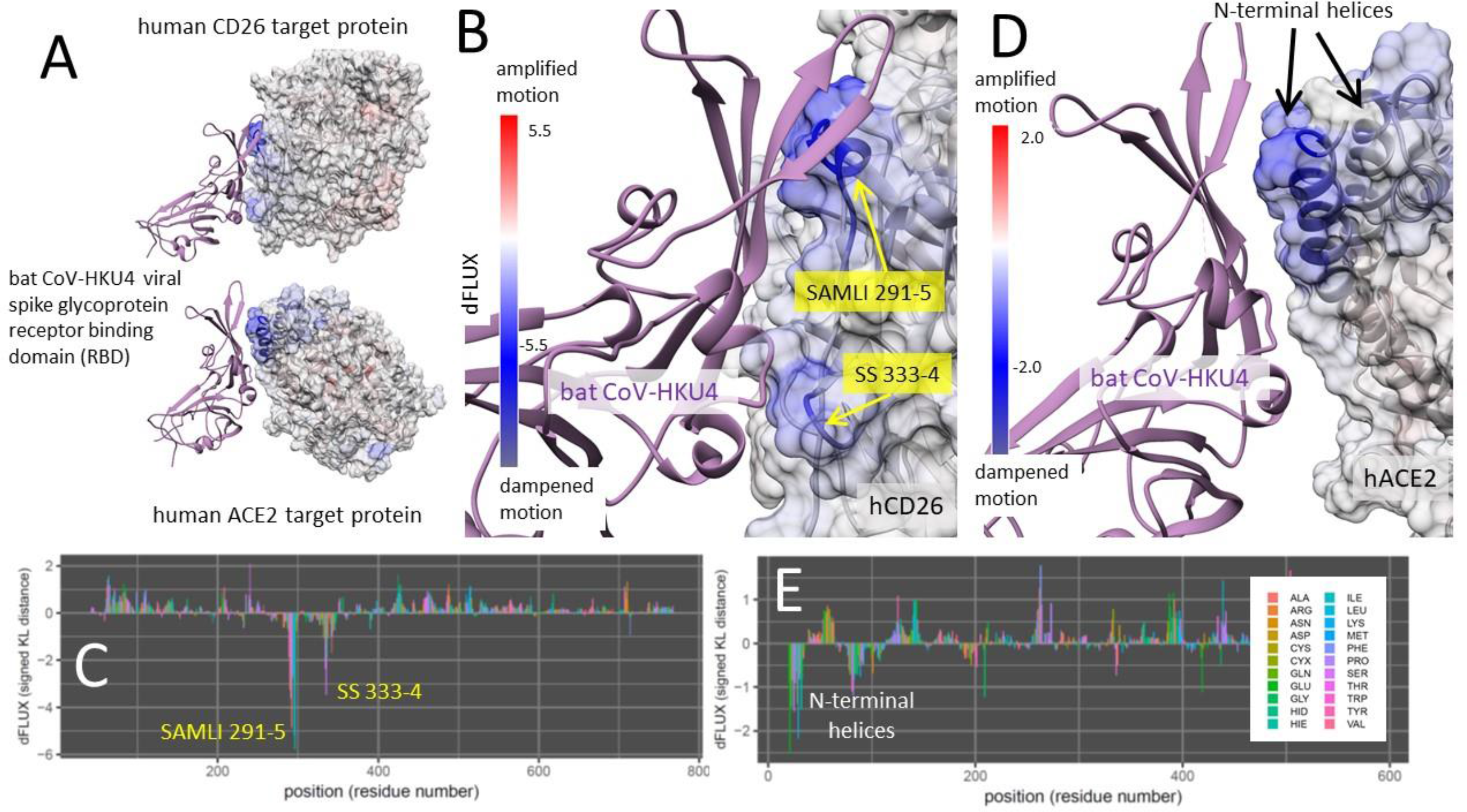
DROIDS binding signature of dampened atom fluctuations in human CD26 and ACE2 receptor proteins upon interaction with the bat progenitor strain batCoV-HKU4 glycoprotein (modeled from PDB: 4qzv and 6m17). Here we show color mapping (A, B, D) and sequence positional plotting (C, E) of dampening of atom motion on the viral RBD-protein target interface in blue for (A-C) its known targeting of CD26 and (A,D-E) its hypothetical targeting of ACE2. The sequence profile of the KL divergence between viral bound and unbound target proteins produces two strong negative peaks indicating key residue binding interactions with the (C) the SAMLI and double serine motifs (shown in yellow) on CD26 and (E) the N-terminal helices on ACE2.

### Summary of the comparative protein dynamic surveys

In comparisons of molecular dynamics simulations between four strains of human-pathogenic coronaviruses, SARS-CoV-2, SARS-CoV-1, hCoV-HKU1, and hCoV-OC43, all four RBDs were associated with statistically significant dampening of molecular motion in the N-terminal helices of human ACE2, indicated by blue color mapping corresponding to a negative Kullback-Leibler (KL) divergence on the ACE2 PDB structure mapped at single amino acid resolution (Figures 2-6). With the exceptions of bat CoV-HKU4 and hCoV-OC43, all RBDs were also associated with dampened molecular motion of ACE2 at residue K353. The absence of a secondary interaction between hCoV-OC43 and ACE2 is further supported by multiple sequence alignment of the RBD loop region proximal to the K353 site, where hCoV-OC43 is the only surveyed strain with a polar uncharged threonine residue in place of a small aliphatic residue (Fig. S6). A significant dampening effect is also present in two novel sites near to K353, the Q325 and 386-AAQPFLL-392 motif, when interacting with SARS-CoV-1 and SARS-CoV-2 (Fig. 5), indicating a transiently dynamic or promiscuous interaction of the viral RBD with these three key ACE2 binding sites. In the recent evolution of the far more transmissible variants (Fig. 6), the stabilization of RBD binding interaction with the N-terminal helices and sites Q325, K353, and 386-AAQPFLL-392 in ACE2 while favoring a more stable K353 interaction highlights an important molecular explanation that linking population transmission rate with the evolution of increased binding specificity in these new variants.

### *In silico* Mutagenesis Study of hACE2 and SARS-CoV-2 Bound Dynamics

To confirm the relative importance of the three promiscuous sites in the binding interaction of coronavirus RBDs to human ACE2, eight total *in silico* targeted mutagenesis studies were performed, on ACE2 and the viral RBD alone (Table 2). K353A, a mutation previously identified to abolish SARS-CoV-1 RBD interaction with ACE2, neutralized dampening of the N-terminal helices during binding with the SARS-CoV-2 RBD (Fig. S7) supporting the observation of decreased infectivity in the mutant (W. Li et al. 2005b). Within the newly identified 386-AAQPFLL-392 motif, a mutation of the nonpolar N-terminal double alanine to the charged polar glutamic acid resulted in an additional strong dampening of molecular motion in the ancestral Q325 site, with a KL divergence of less than −3 between the unbound and bound states. Mutating the C-terminal FLL residues to EEE in the 386-AAQPFLL-392 motif resulted in no overall change to the binding characteristics between the SARS-CoV-2 RBD and ACE2. Upon mutating V739 in ACE2 to hydrophilic glutamate, the N-terminal helix region and 386-AAQPFLL-392 motif showed no significant change in molecular motion between bound and unbound states (Fig. S8). This same pattern of binding characteristics could be seen in the mutation of V185 to glutamate in the loop region of the SARS-CoV-2 RBD, proximal to ACE2 during binding (Fig. S9A). Mutating V185 to leucine in the RBD resulted in the same binding characteristics, resulting in a pattern of dampening in molecular motion that was the same as seen in SARS-CoV-1 (Fig. S9B). All surveyed mutations maintained significant dampening in K353 of ACE2 during binding. The mutation of V739 in ACE2 to leucine resulted in dampening of Q325 in the interaction, as is seen in SARS-CoV-1, and a lack of dampening in the AAQPFLL motif as seen in the SARS-CoV-2 wild type. However, mutating V739 in ACE2 to glutamate resulted in pronounced dampening only at the N-terminal helices and K353. Upon mutating polar K408 in the SARS-CoV-2 RBD to nonpolar alanine, the Q325 interaction disappeared and only interactions with N-terminal helices, K353, and 382-AAQPFLL-392 remained (Fig. S10).

### Machine learning-based detection of functionally conserved ACE2 binding dynamics at sites shared between human/bat *Betacoronavirus* orthologs

The maxDemon algorithm detected significant regional canonical correlations in the machine learning classification performance profiles of bat/human and SARS/SARS variant orthologs, indicating regions of functionally conserved dynamics that was recognizable from thermal noise and shared between the molecular dynamics modeling of even distant orthologs (Fig 9A-C). The most distant evolutionary comparison of bat CoV-HKU4 to SARS-CoV-2 identified functionally conserved dynamics at 4 highly localized sites including the region surrounding K353 (Fig 9A). The comparison of the bat CoV-HKU4 to human hCoV-HKU1 identified functionally conserved dynamics at 2 of the 3 N-terminal helices, as well as two of the same highly localized regions observed in the comparison of the bat progenitor to SARS-CoV-2 (Fig 9B). However, the K353 site was not conserved in the comparison with the human endemic strain. In the closest evolutionary comparison of the dynamics of SARS-CoV-2 and its B.1.1.7 variant, we observed larger regions of functionally conserved dynamics that includes all the regions observed in the more distant comparisons (Fig 9C). These regions included all the N-terminal helices, as well as the regions connecting the key sites at K353 and Q325. The protein dynamics from position 150-200 and 280 to 308 were also significantly conserved. The consistency of the detection algorithm across these different simulations, as well as the decrease in functionally conserved dynamics with increased evolutionary distance, very reminiscent of DNA sequence-based methods, was highly encouraging despite the ‘black box’ nature of machine learning approaches to complex identification problems such as those posed when trying to compare molecular dynamic states of protein function.

**Figure 9.**
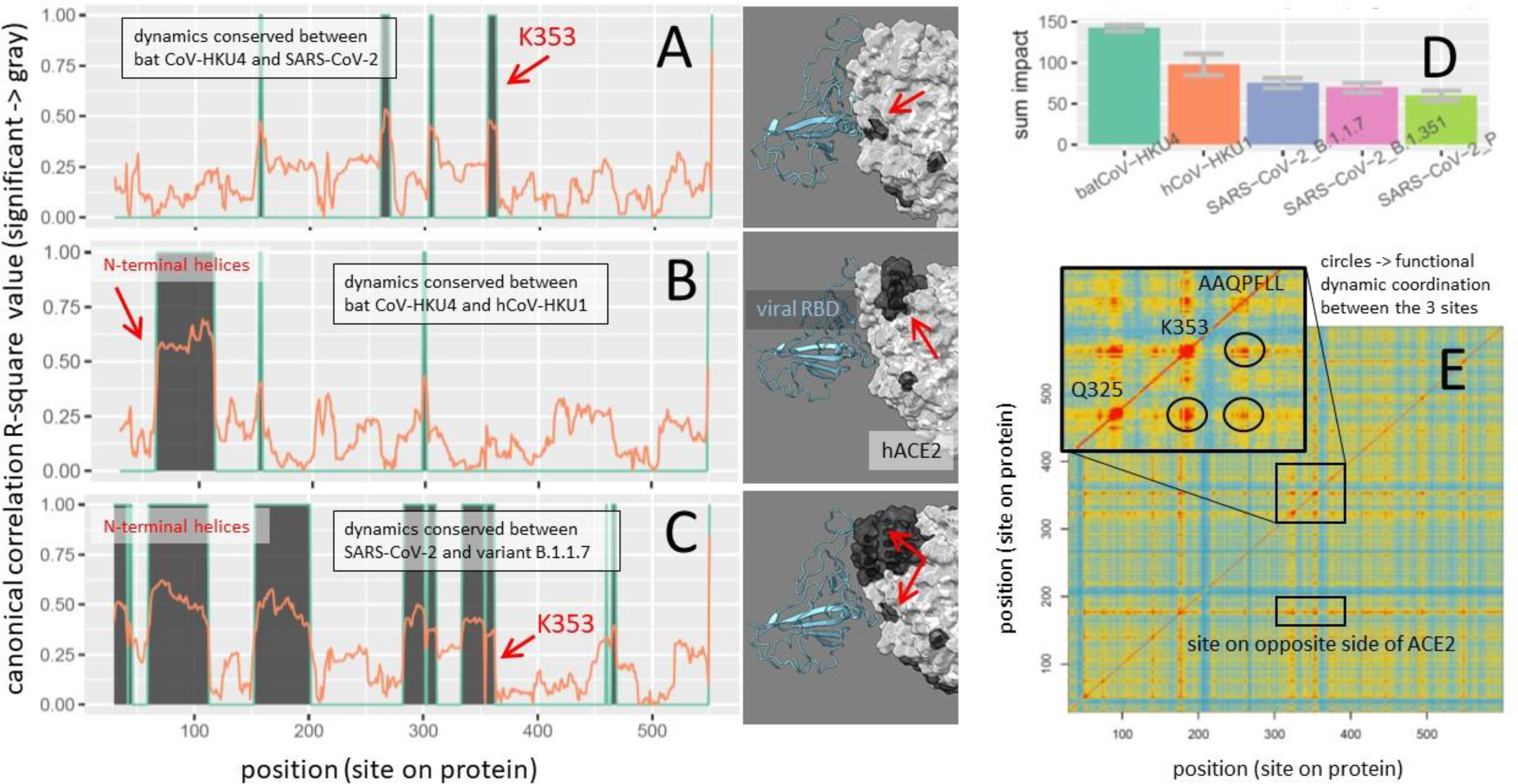
Functionally conserved and coordinated binding dynamics. (A-C) Maps of evolutionary conserved functional binding dynamics (shown in dark gray) of viral RBD on both ACE2 sequence and structure. Three different evolutionary distances were considered including the most distant comparison between SARS-CoV-2 and (A) the hypothetical bat CoV-HKU4 progenitor, (B) the human endemic strain hCoV-HKU1, and (C) the UK variant, SARS-CoV-2 B.1.1.7. (D) These evolutionary distances are also represented as the total sum of the mutational impacts on conserved dynamics. (E) The coordination of the functionally conserved binding dynamics of ACE2 upon interactions with SARS-CoV-2 receptor binding domain is shown as a matrix with red/blue indicating high/low pairwise coordination (i.e. mutual information in learning classifications) between sites. The inset highlights the high degree of coordination occurring between K353, Q325 and the AAQFPLL motif in their dynamic interactions with the viral N501 site.

### Information theoretics for comparing variants and detecting functional coordination in ACE2 sites of interest

The total impact of the genetic differences among the *Betacoronovirus* strains (i.e. relative entropy of correlation) upon the functionally conserved dynamics detected by the machine learning model followed a quite predictable pattern, with more evolutionary distant comparisons exhibiting more total impact on dynamics (Figure 9D). The mutual information in learning classifications between sites also captured the predicted coordination in functional binding that we expected to see in the dynamics at K353, Q325 and 382-AAQPFLL-392 (Figure 9E). Not only were the functional dynamics coordinated across amino acid sites near each of these binding events (red regions on the diagonal of the heatmap) but they were coordinated across these three sites as well (at circled intersections off the diagonal). A two-dimensional representation of the machine learning pre-classifications and discriminant learning space for the training data at each of the three key sites and a site on the N-terminal helices is also shown (Fig 10). Here, we can observe that the K353 site clearly has the highest signal to noise ratio regarding the functional binding of SARS-CoV-2 RBD and human ACE2.

**Figure 10.**
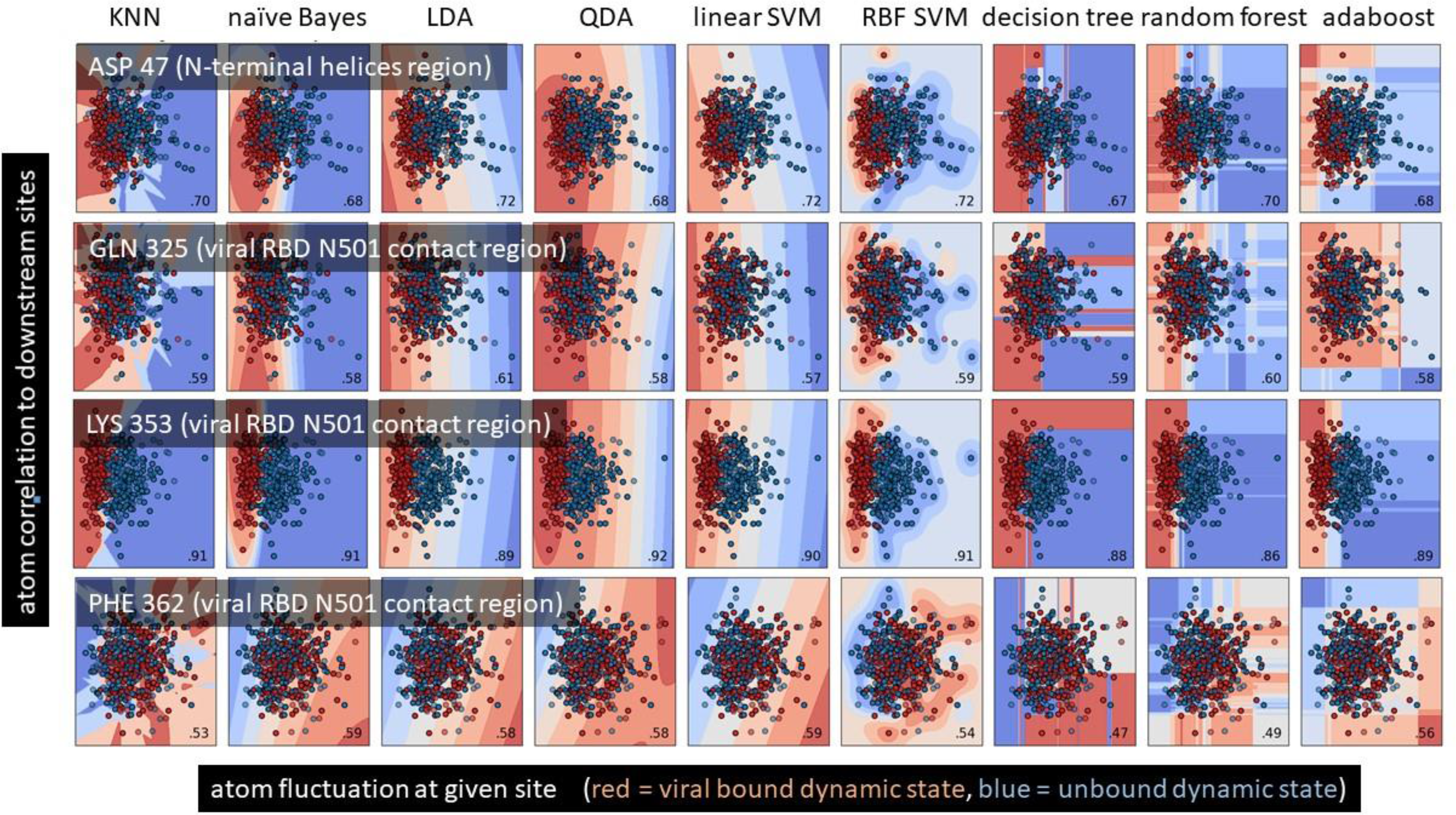
Feature vector and learning classification spaces for key binding sites on ACE2. The machine learning training data and the optimized learning classification spaces is shown at four key residue sites on the binding interface of SARS-CoV-2 and ACE2. The X axis is the atom fluctuation at the given site and the Y axis is the correlations of the fluctuations with nearby downstream sites. The data points represent individual 50 frame time slices in the simulations with dark red indicating those time slices pre-classified to the viral bound state of ACE2 and dark blue points indicating those time slices pre-classified to the unbound state of ACE2. The colors of the background indicate the probability state of the machine learning classification model after training on the data points. The average correct classification of the data set is shown in the lower right corner of each plot.

**Figure 11.**
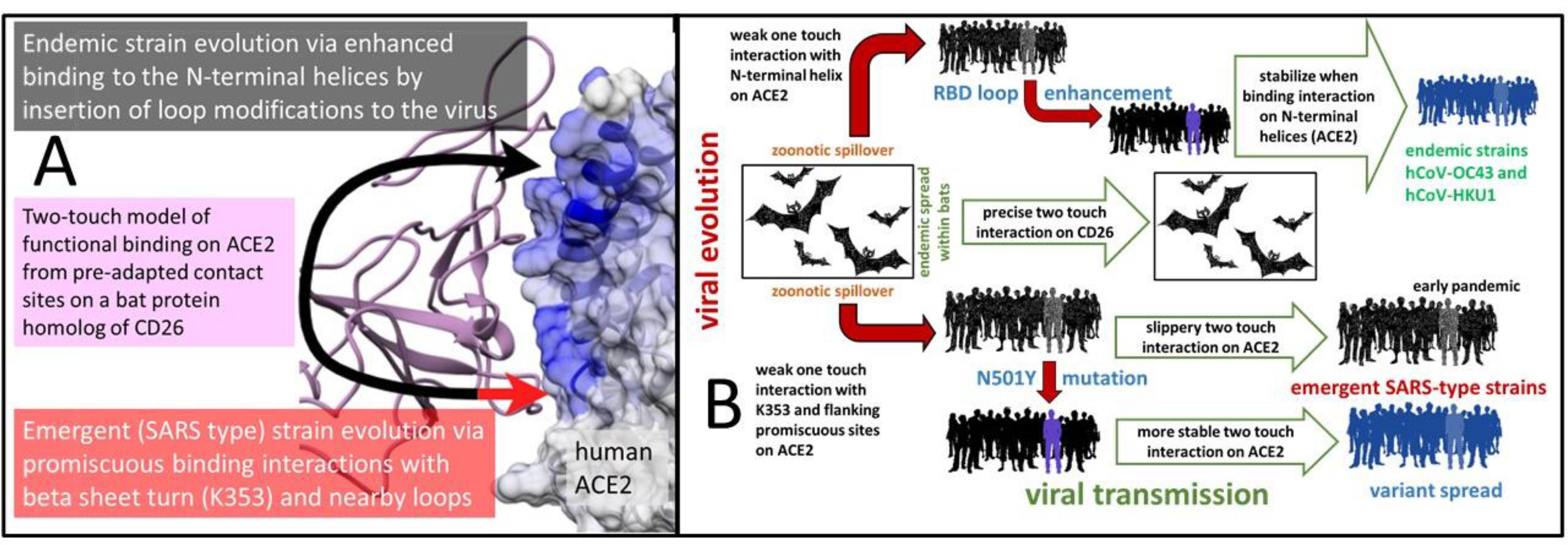
A functional evolutionary model of SARS-type and endemic viral protein binding interactions during zoonotic spillover. (A) The two-touch interaction model consists of a combination of evolutionary conserved ACE2 interactions at the N-terminal helices likely preadapted from its interaction with the SAMLI motif on CD26 (shown with black arrow) and a promiscuous, dynamically complex interaction with K353 and nearby sites at Q325 and AAQPFLL 386-392 (shown with red arrow), likely preadapted from its interaction with SS 333-334 on CD26. (B) Given the conserved nature of viral receptor binding across the diversity of strains we surveyed, we hypothesize these regions of the virus are most likely responsible for past and present instances of zoonotic spillover to humans. The more pathogenic SARS-like outbreaks associated with ACE2 seem to develop transient dynamics of N501 that involve promiscuous interactions at three co-located sites mentioned above. This may create a temporary ‘slippery two touch interaction with the ACE2 target that stabilizes of evolutionary time. In the much more transmissible new variant strains, (B.1.1.7, B.1.351, and P.1), the promiscuous interactions are mostly lost, favoring a much more stable binding interaction with K353 alone. In both endemic strains, we observe the evolution of enhanced binding to the N-terminal helices of ACE2 through the evolution of enhanced loop structure on the corresponding interface of the viral receptor binding domain.

### A model of the functional evolution of ACE2 binding interactions during zoonotic spillover

Based on our results comparing regions of functionally conserved dampened molecular motion in ACE2 and CD26 target proteins across zoonotic, emergent, and endemic *Betacoronavirus* strains, we propose a model of protein interaction for coronavirus RBD evolution and human specificity highlighting the role of the two distinct touch points between the viral RBD and the human target proteins (Fig 11A). The black arrowhead in the model signifies a relatively conserved region of sustained interaction between the coronavirus RBDs and the N-terminal helix region of ACE2. The red arrowhead indicates more promiscuous interactions with Q325 and K353, present individually across SARS-CoV-2 as well as SARS-CoV-1, MERS, and HKU1, as well as the newly identified 386-AAQPFLL-392 motif prominent mainly in ACE2 interactions with the wild-type SARS-type spike protein RBD. This two-touch model highlights a common interaction point of the viral RBD with the N-terminal helices of ACE2 that is conserved in all known strains of potentially human-interacting *Betacoronavirus*, demonstrated by the universal dampening of ACE2 molecular motions upon viral binding. This interaction corresponds structurally with the viral bat progenitor strain RBD interaction with the SAMLI 291-5 site in CD26. The second and more complex dynamic touch point in the model involves the contact of the viral RBD N501 with the previously described contact sites K353 and Q325, as well as the nearby novel site of interaction at the AAQPFLL motif. This second interaction corresponds with a strong and precise binding interaction of the viral bat progenitor strain RBD with SS 333-4 in CD26. This promiscuous interaction site seems considerably apparent in the two pathogenic SARS-type strains, suggesting that the binding interactions during zoonotic spillover may follow a common functional evolutionary path in SARS-type emergent outbreaks (Fig 11B), perhaps utilizing a preadapted interactions with the SAMLI and SS motifs in CD26 in bat reservoir populations to establish loose and promiscuous binding with the ACE2 N-terminal helices and the loop regions near the beta sheet turn at K353. In SARS-CoV-2, this has most recently evolved to higher specificity for ACE2 particularly through the strengthening of the ACE2 K353 and the N-terminal interactions. This appears quite different from the evolutionary path of the binding interactions in human endemic strains like hCoV-OC43 (Fig 11B), where an insertion event appears to have enhanced the RBD loop structures near the N-terminal helices of ACE2 and subsequently strengthen binding interactions in this region.

## DISCUSSION

Here we conducted machine-learning assisted statistical comparisons of hundreds of replicate sets of nanosecond scale GPU accelerated molecular dynamics simulations on viral bound and unbound target proteins to survey the evolutionarily conserved functional similarity as well as the comparative differences between various emergent, endemic, and potentially zoonotic *Betacoronavirus* strains in humans. We present evidence that strongly suggests a common route of functional molecular evolution occurring at the binding interface of the viral receptor binding domain and their primary protein targets ACE2 and CD26. We propose a two-touch contact model of viral evolution that may be evolving similarly during SARS-type *Betacoronavirus* zoonotic spillover events from either *Tylonycteris* (bamboo bat) or *Rhinolophus* (horseshoe bat) to human populations. We present computationally derived signatures of receptor binding domain interactions with target human proteins, in the bat progenitor strain batCoV-HKU4 and five human strains, the emergent MERS-CoV, SARS-CoV-1 and SARS-CoV-2 strains and the endemic hCoV-OC43 and hCoV-HKU1 strains. We use novel machine-learning based approach to detect functionally conserved ACE2 binding dynamics among viral ortholog comparisons. Our studies of the biophysics of viral binding at single amino acid site resolution demonstrate that the coronavirus spike glycoprotein receptor binding domain (RBD) has a strong, static, and well-defined binding interaction occurring in two distinct regions, or “touch points” of human CD26 (also likely the CD26 ortholog in bats as well). In its interactions with the human ACE2 target protein, the spike glycoprotein RBD appears pre-adapted through its binding interactions involving two of these sites on CD26 (SAMLI 291-5 and SS 333-4) to allow broad and somewhat promiscuous binding interactions with the N-terminal helical region of ACE2 as well as regions connecting K353, Q325 and 382-AAQPFLL-392.

We find that the functional binding dynamics of these two pre-adapted contact regions on ACE2 have been conserved differently between the bat progenitor and the human endemic and recent SARS-type emergent strains. The human endemic strains hCoV-OC43 and hCoV-HKU1 appear to have evolved towards enhanced stability on ACE2 via strong broad interaction with the ACE2 N-terminal helices that is facilitated by the evolution of enhanced loop structures on the viral RBD that markedly stabilized upon binding the interface with N-terminal helices on ACE2. The slightly more pathogenic human strain hCoV-HKU1 also shows shares this feature with hCoV-OC43, but still also shows weak interactions with K353. Thus, while the binding of the ACE2 N-terminal helices appears common within all of our *Betacoronavirus* strain dynamics models, this interaction appears especially conserved in the functional evolution of past endemic human strains from the bat progenitor. However, in the more emergent SARS-type strains, we observe evolutionary conserved binding dynamics mainly at and near the K353 site on ACE2, where we also observe a high degree of mutual information (i.e. coordination of functional dynamic states) between this site and Q325 and 382-AAQPFLL-392. This transient viral RBD interaction with multiple sites on ACE2 in emergent SARS-type strains suggests that the interaction with K353 reported by (Yan et al. 2020), is not as stable in the early evolution of SARS-CoV-2 binding as their initial structural analysis might imply. We hypothesize that the binding interactions at these three sites are perhaps more evolvable in SARS-type strains than the interactions at the N-terminal helices. The recent appearance of the N501Y in the highly transmissible variants of SARS-CoV-2 appears to further support the functional evolution of more stable binding at this same location on the ACE2 target protein. When we characterized the effect of the mutation on the binding dynamics in this region, we found that this one mutation has greatly stabilized the Y501 interaction with the ACE2 K353 site and reduced the transient and promiscuous interactions with the other two flanking binding sites. Thus, the stabilization the dynamics at of this point of contact has enhanced the binding, and by extension perhaps also the transmissibility of the SARS-CoV-2 virus, all the while bringing it much closer to the molecular binding profile of the bat progenitor strain batCoV-HKU4 with its primary target protein, CD26. It has been recently hypothesized and demonstrated that promiscuity in protein-protein interactions is related to a lack of protein stability (Cohen-Khait et al. 2017) and it would also appear, through our work on this system, that a lack of protein stability may also be related to the molecular facilitation of zoonotic spillovers of certain viral pathogens as well. We believe the complex dynamic interaction at the second ‘slippery or transient touch point’ in our two-touch model may represent the molecular manifestation of a lack of binding specificity that may characterize many viral binding interactions during emergent SARS-type outbreaks when the co-evolution between a viral binding domain and a potential host target receptor is still historically very recent. Our simulations suggest several alternative paths of viral evolution in emergent versus endemic strains that have both favored a more precise and specific targeting of human ACE2, which may also be associated with enhanced transmissibility as well.

While our approach offers the considerable advantage of combining a comparative statistical method and a physics-based modeling approach towards addressing functional molecular evolution, it is not without some pitfalls. Some potential limitations of MD simulations as a probative method for functional molecular evolution are its many implicit simplifying computational assumptions, its complex and inherently stochastic nature, and high computational expense (i.e. due to femtosecond time steps). Specifically, computational sampling of even the accelerated MD method employed here has runtime limitations, and even on modern graphics cards our simulations can typically have a cumulative runtime of several weeks to generate the proper statistical replication to compare physical time frames of only several hundreds of nanoseconds. In addition, MD simulations always involve some simplification of the biophysics within the system being studied as it invariably ignores atomic charge regulation, bond motions in the solvent, charge screening during interaction, and other macromolecular crowding effects. Insight into long-term microsecond to millisecond dynamics in large explicit solvent systems are still limited by currently available hardware, even when advanced algorithms for accelerating MD simulations are used. Glycosylation is another aspect of coronavirus spike protein biology that is not fully captured by our MD simulations, mainly due to lack of glycosylation in the functional binding interface of most of our key starting structures. While glycosylation probably does affect the flexibility of ACE2 (Barros et al. 2021), we would argue that while these post-translational modifications can have in immediate impacts on dynamics, their probable lack of genetic heritability justifies our ignoring their role during longer term viral evolution outside of specific outbreaks.

Despite these limitations, we conclude that our identification of additional key residues in the binding interaction between the SARS-CoV-2 RBD and human ACE2 receptor, as well as our evolutionary exploration of the two-touch model of RBD evolution provide a conceptual framework for future functional mutagenesis studies of this system. This will be especially important for understanding the functional evolution of transmissibility in new SARS-CoV-2 variants, recently responsible for the large community lockdowns during the COVID-19 pandemic and recently implicated in reduced vaccine efficacy (Planas et al. 2021). Future surveys of *Betacoronavirus* circulating in past, present, and future human populations as well as molecular and clinical investigations of SARS-CoV-2 infection will likely continue to be further informed by the model interpretation of binding interaction that we present in this study. In future studies of computational models derived from potential zoonotic coronavirus strains, our binding model can lend greater interpretability to observations regarding evolutionary diversity in coronaviruses infecting reservoir species like bats, birds, and small mammals. Finally, as recent work has called out the importance of specific molecular motions in forming novel therapeutic targets for intervention in emerging zoonotic spillover events (Pierri 2020), tools for highlighting functionally conserved dynamics of conformational alterations of the interaction host and pathogen proteins will prove a valuable addition to the arsenal of modeling approaches available for drug development.

## DATA AVAILABILITY STATEMENT

Structural models, images, movies and data sets for the post-processed molecular dynamics presented in this work are deposited at Zenodo CERN under the DOI 10.5281/zenodo.4702398 https://zenodo.org/record/4702398

A movie file comparing SARS-CoV-2 RBD interaction with ACE2, in both wildtype and N501Y mutant strains is also available at https://youtu.be/ptn1_BBJi70

## SUPPORTING MATERIAL

The main repository for DROIDS 4.2 and maxDemon 2.0 can be found at the Website + GitHub repository link below. Please follow the link to “Releases” and download the latest release at https://gbabbitt.github.io/DROIDS-4.0-comparative-protein-dynamics/

## AUTHOR CONTRIBUTIONS

PR, FC and GAB designed and executed the biological study. GAB designed the statistical methods and wrote computer code to implement the methods. PR, GAB, FC, AOH, and MLL interpreted the results and wrote the manuscript.

## ACKNOWLEDGEMENTS

We acknowledge funding support from the National Science Foundation (NSF Award Number 2029885) and the National Institutes of Health (NIH grant GM116102). We also acknowledge the Nvidia Corporation for a hardware support grant.

## Notes

https://zenodo.org/record/4702398

### Competing Interest Statement

The authors have declared no competing interest.

### Summary of Updates

This revision adds new data pertinent to the new N501Y mutations of SARS-CoV-2 in the highly transmissible UK and South African/Brazilian variants. It also provides a machine learning analysis of the evolution of emergent and endemic Betacoronavirus strains

