## Supplemental Figures for "Functional binding dynamics relevant to the evolution of zoonotic spillovers in endemic and emergent *Betacoronavirus* strains"

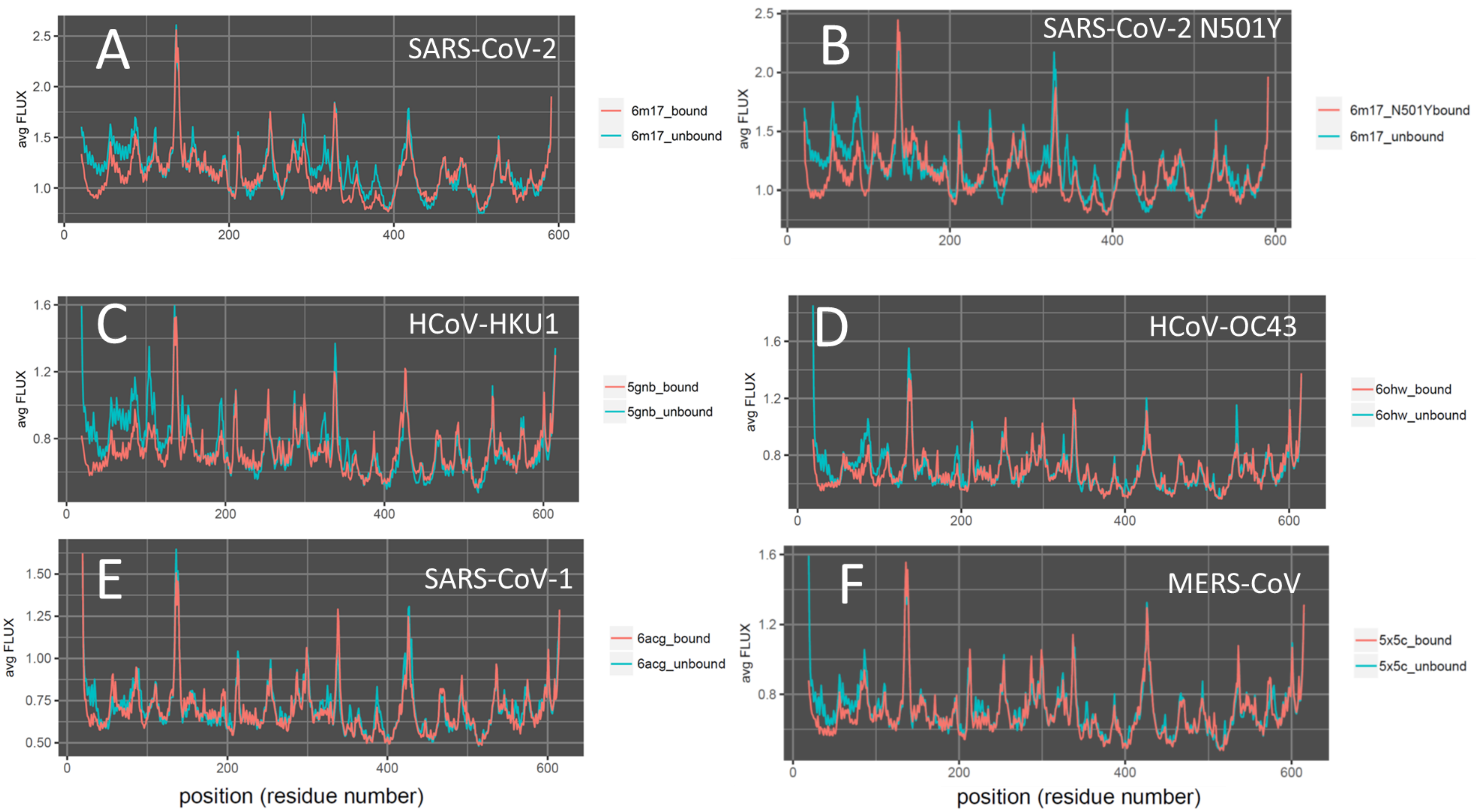


**Supplemental Figure S1.** Amino-acid site-wise average root mean square fluctuation profiles for the viral bound and unbound ACE2 targets in this study.

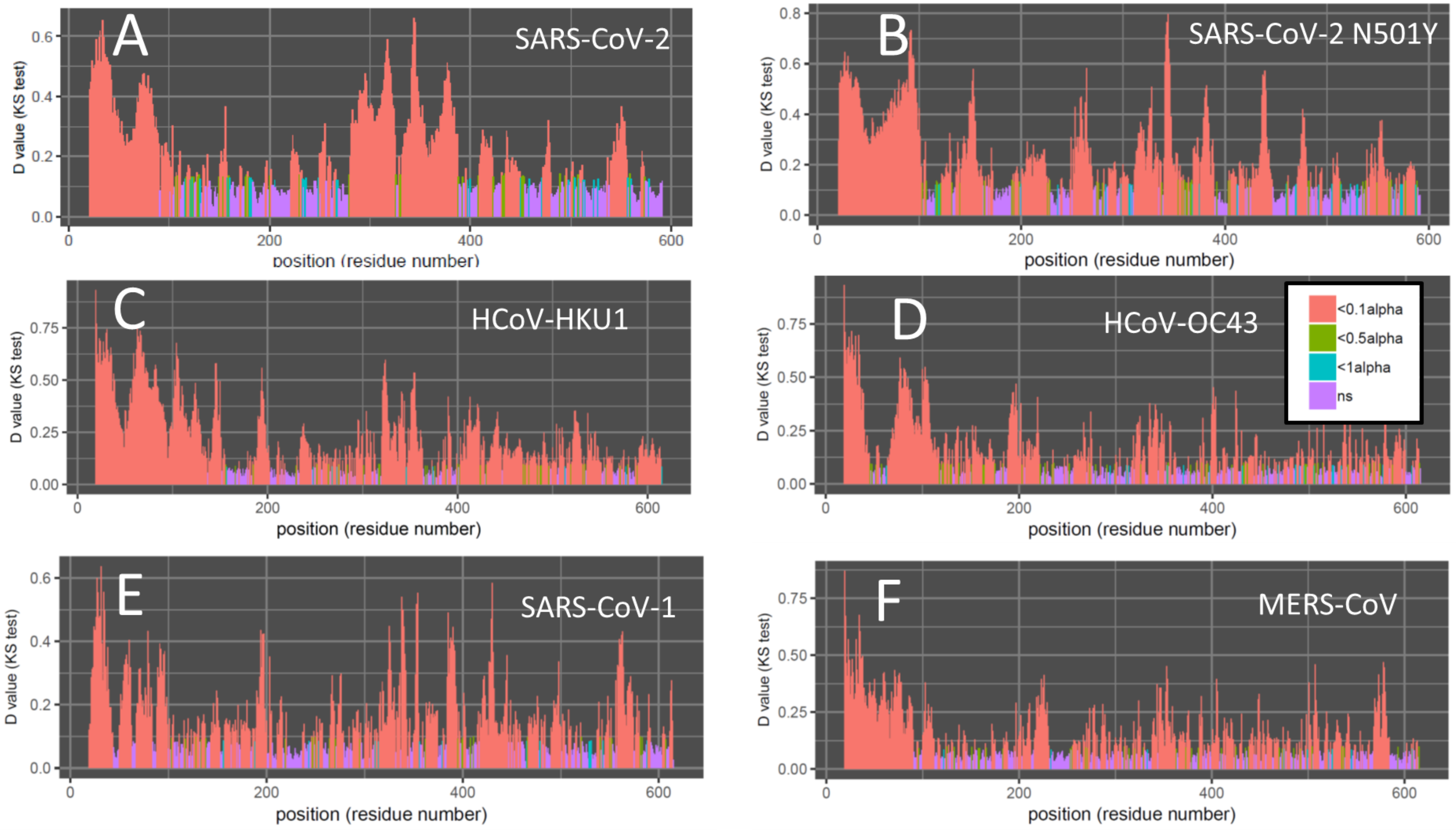


**Supplemental Figure S2.** Amino-acid site-wise significance tests for the root mean square fluctuation differences between the viral bound and unbound ACE2 targets in this study. The test of significance is a two sample Kolmogorov-Smirnov test with a Benjamini Hochberg p-value correction to account for the number of sites. The plots show the D value (i.e. test statistic) color coded by p-value with orange indicating $p < 0.005$, green indicating $p < 0.025$, aqua indicating $p < 0.05$ and violet indicating $p > 0.05$. Here the p-value of the two sample KS test indicates the probability of the viral bound and unbound ACE2 protein dynamics are drawn from the same population.

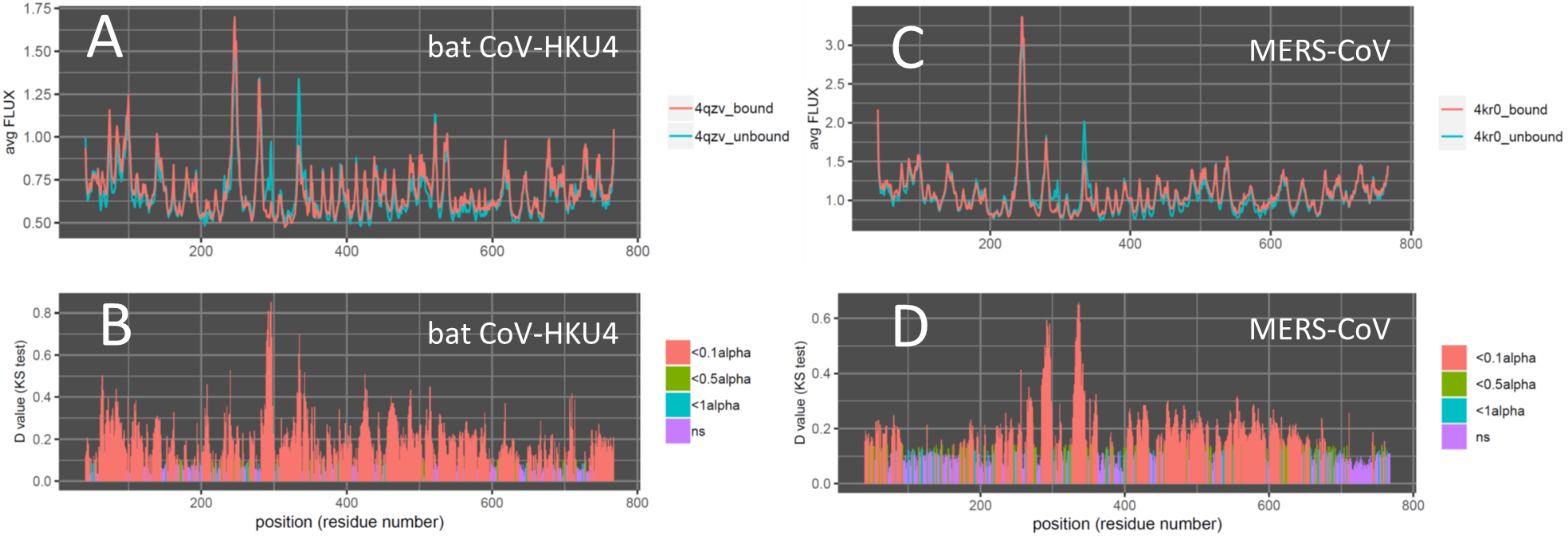


**Supplemental Figure S3.** Amino-acid site-wise fluctuation profiles and significance tests for the root mean square fluctuation differences between the viral bound and unbound CD26 targets in this study. The test of significance is a two sample Kolmogorov-Smirnov test with a Benjamini Hochberg p-value correction to account for the number of sites. The plots show the D value (i.e. test statistic) color coded by p-value with orange indicating $p < 0.005$, green indicating $p < 0.025$, aqua indicating $p < 0.05$ and violet indicating $p > 0.05$. Here the p-value of the two sample KS test indicates the probability of the viral bound and unbound ACE2 protein dynamics are drawn from the same population.

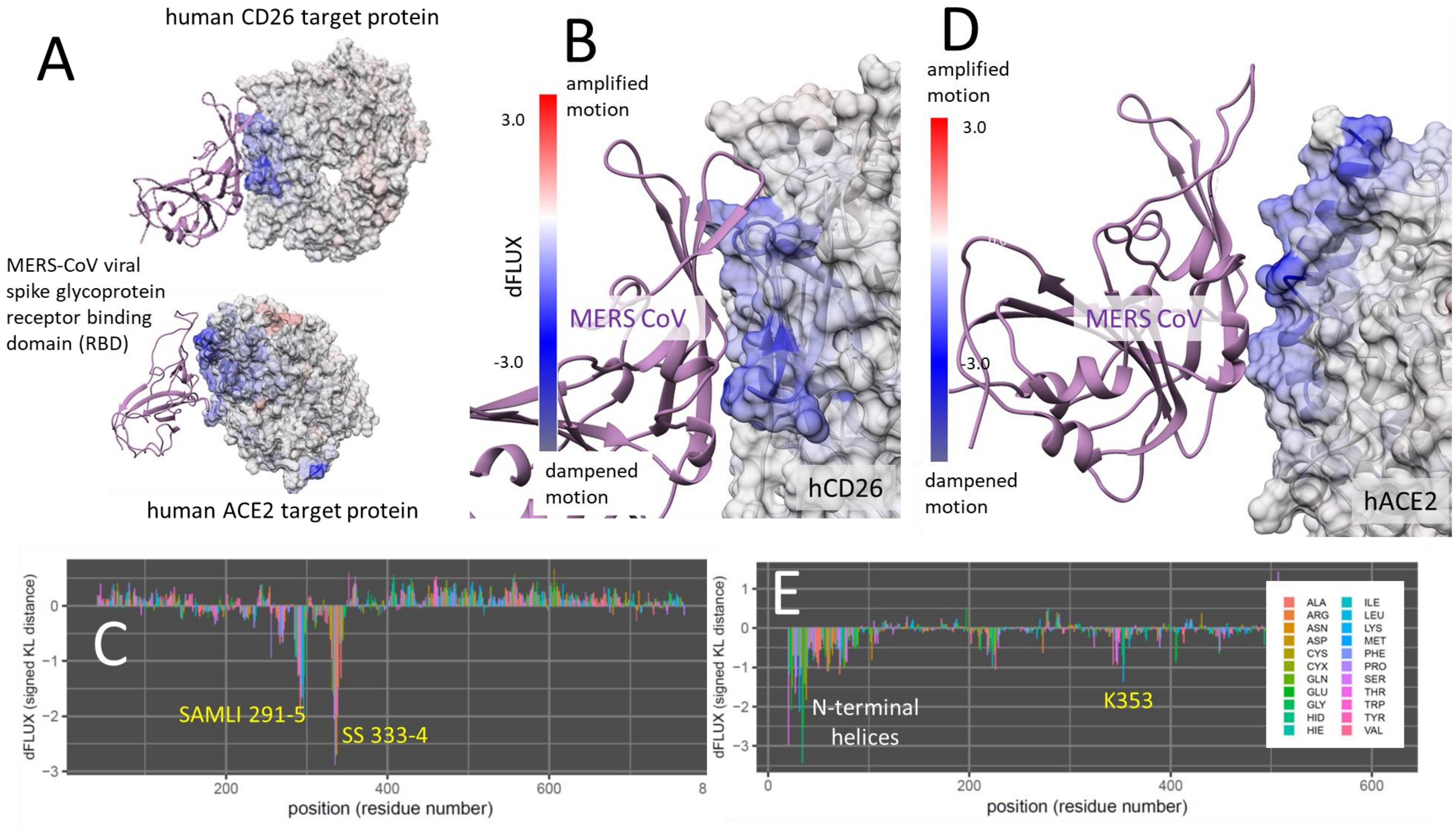


**Supplemental Figure S4. DROIDS binding signature of dampened atom fluctuations in human ACE2 receptor proteins upon interaction with the past human outbreak strain MERS-CoV spike glycoprotein (modeled from PDB: 4kr0, 5x5c, and 6m17).** Here we show color mapping (A, B, D) and sequence positional plotting (C, E) of dampening of atom motion on the viral RBD-protein target interface in blue for (A-C) the targeting of CD26 by the MERS-CoV and (D-E) the hypothetical targeting of ACE2 by MERS-CoV. The sequence profile of the KL divergence between viral bound and unbound target proteins produces strong negative peaks indicating key residue binding interactions with (C) the SAMLI 291-5 and SS333-4 motifs on CD26 and € with the N-terminal helices on ACE2. This is only a very weak interaction with K353 (yellow) on the ACE2.

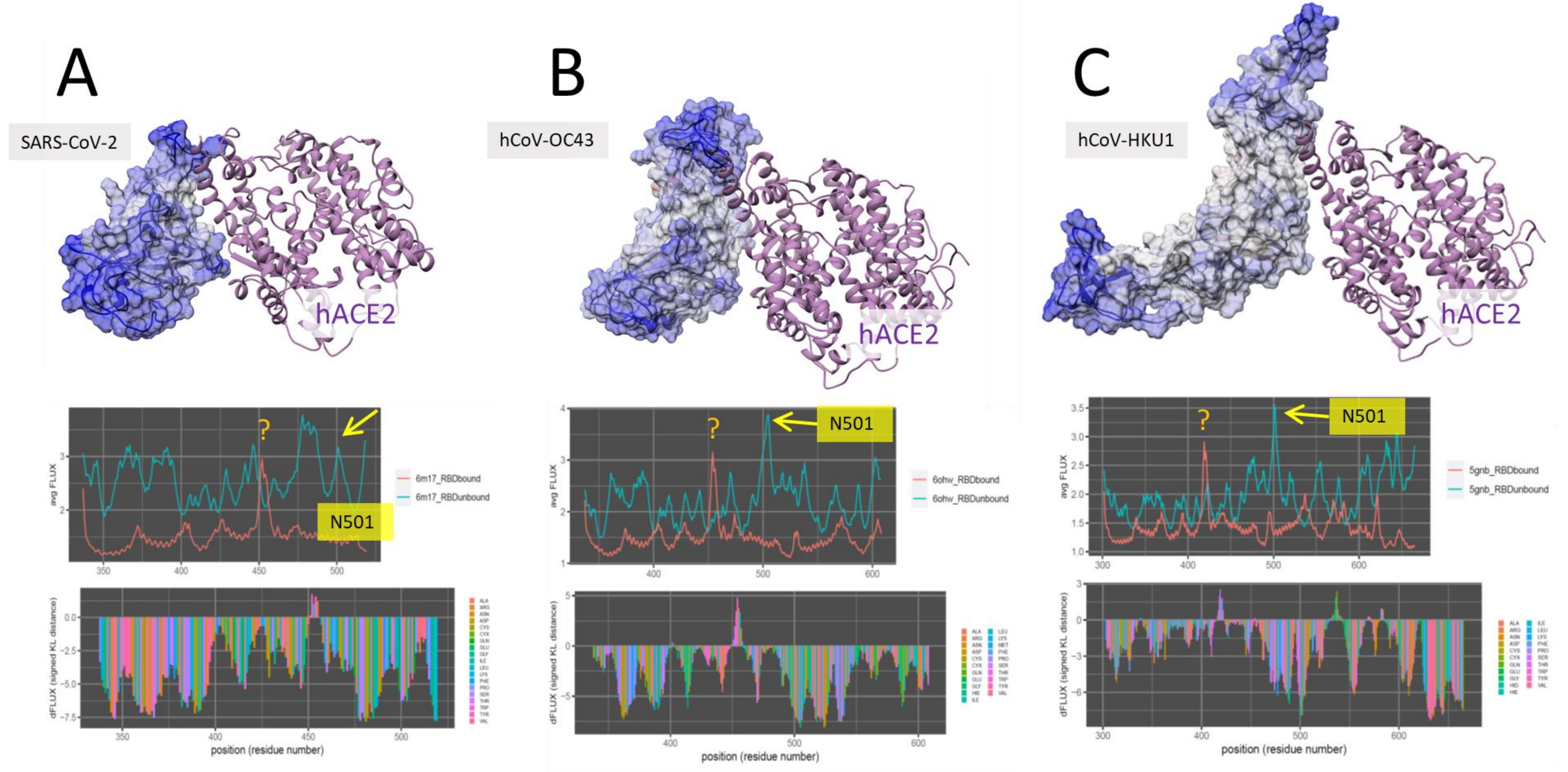


**Supplemental Figure S5.** DROIDS binding signature of dampened atom fluctuations in emergent and endemic viral receptor binding domains upon interaction with human ACE2.

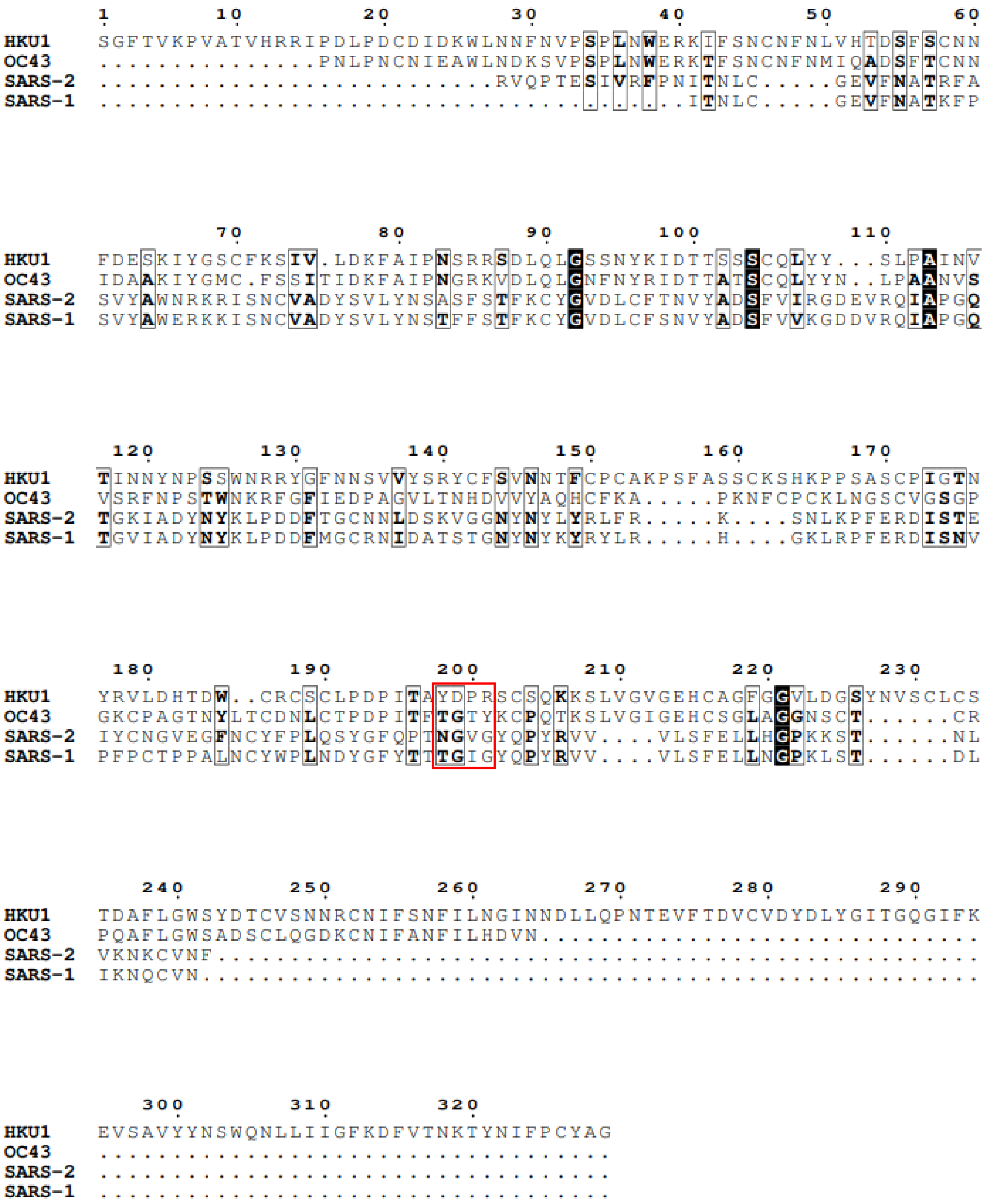


**Supplementary Figure S6.** MSA of human coronavirus RBD amino acid sequences derived from PDB structures. Boxed in red is the loop region proximal to the second ACE2 touch point, where differences in chemical properties between OC43 and other coronavirus strains can be seen particularly at position 200.

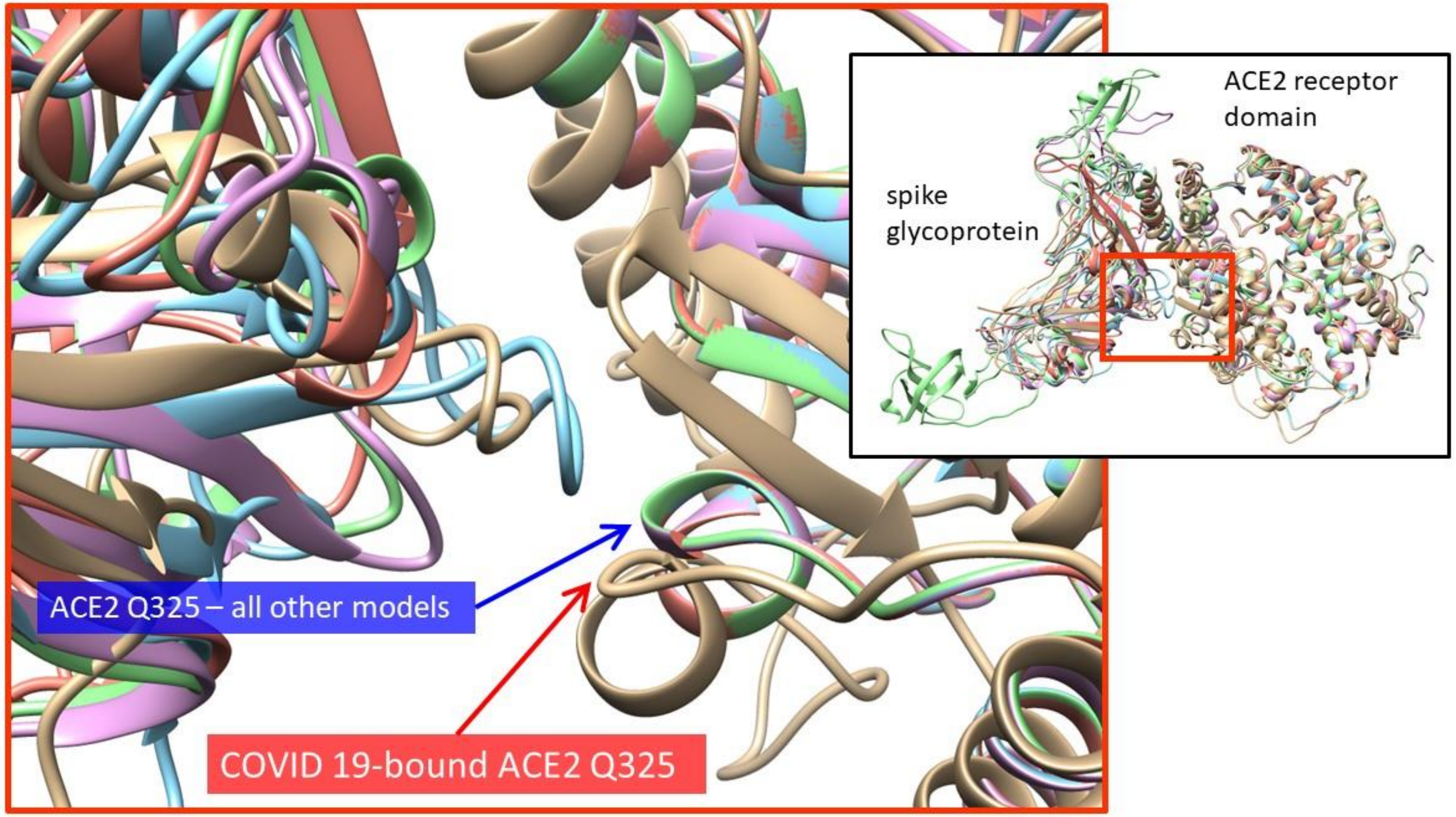


**Supplemental Figure S7.** Structural alignment of the five outbreak models used in our study (tan=SARS-CoV-2(COVID19), aqua=SARS-CoV-1(classic SARS), red=MERS-CoV, lavender=HCov-OC43 and green=HCoV-HKU1).

Close-up image highlights the structural difference at Q325 caused by COVID 19 from the four other models.

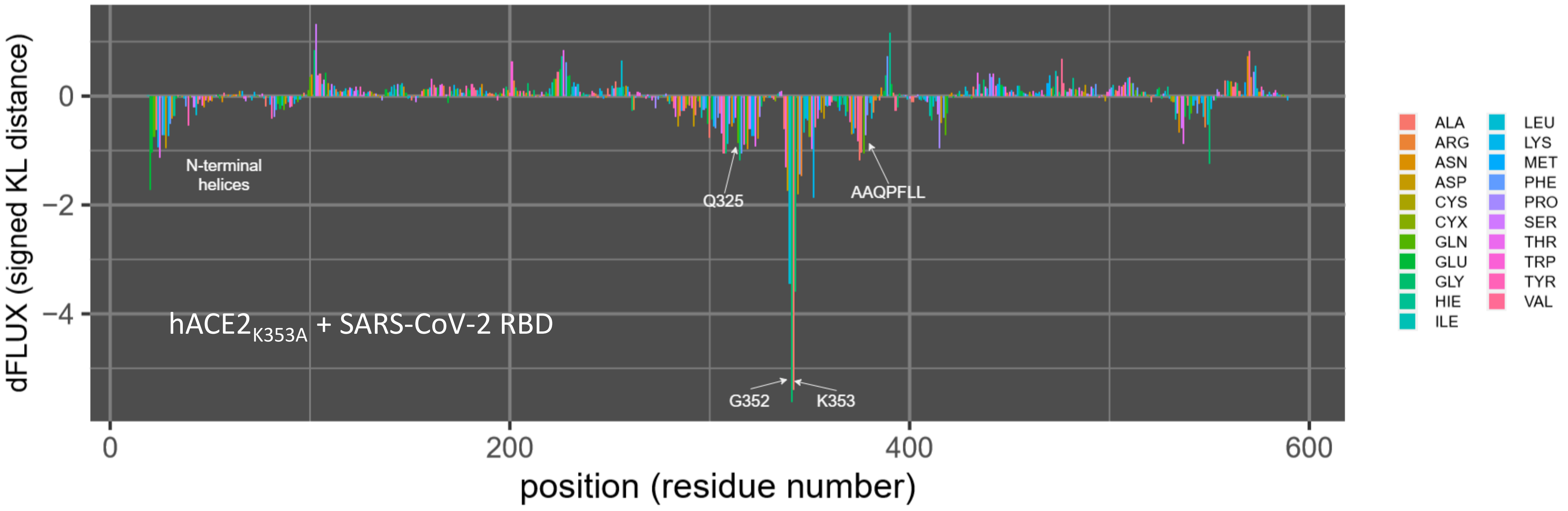


**Supplemental Figure S8.** Per-residue rmsf of SARS-CoV-2 RBD in complex with human ACE2 mutated at the K353 position in silico. Mutagenesis was performed with the swapaa command in Chimera v. 1.13 and the structure was minimized with 2000 steps of steepest descent. Residues of interest in comparison to the wild-type SARS-CoV-2 RBD/ACE2 complex are labeled. This mutagenesis study was conducted to validate the ability of the DROIDS 3.0 molecular dynamics tool to corroborate RBD/ACE2 interaction-discouraging amino acid substitution K353A in SARS-CoV-1 [34].

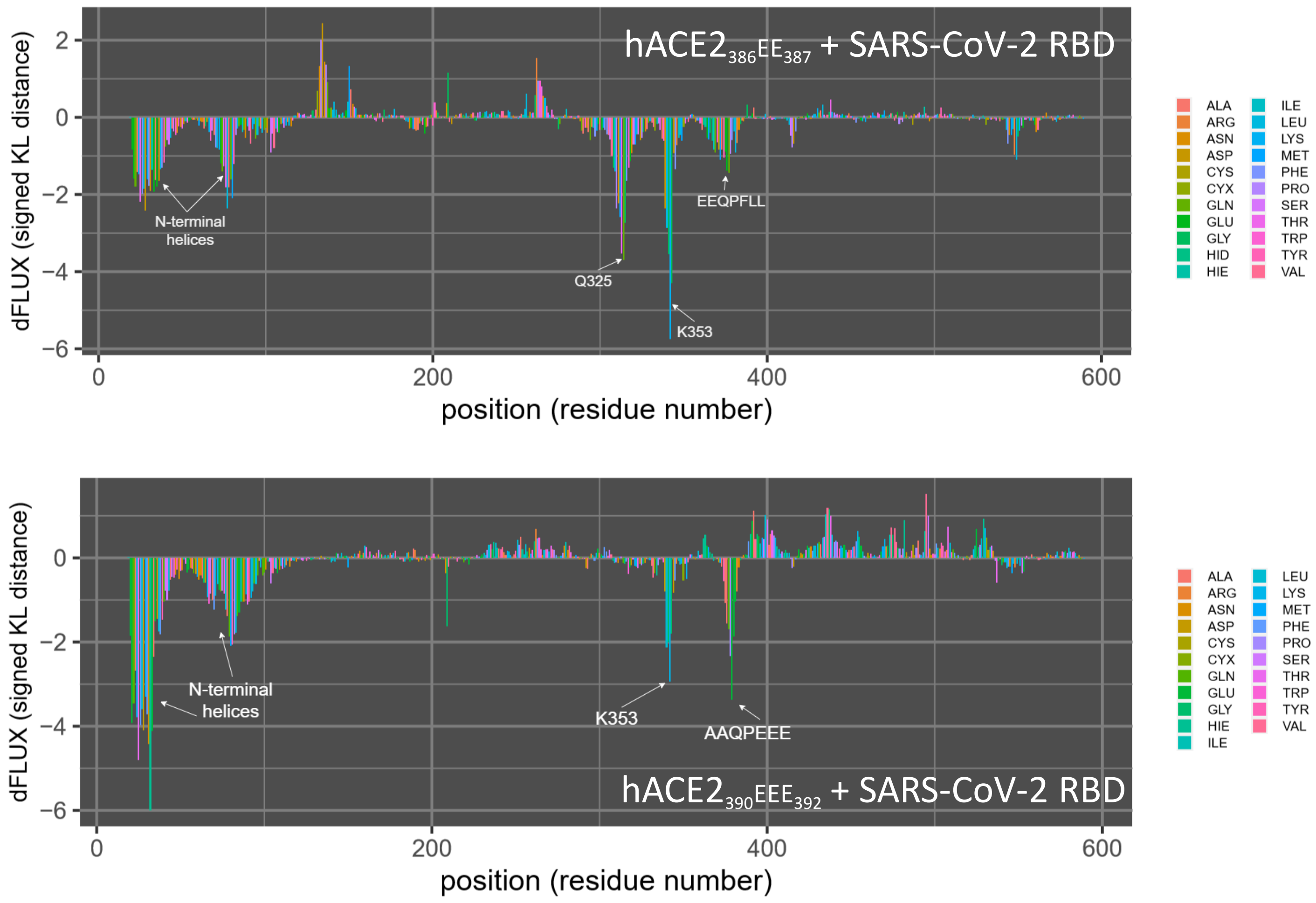


**Supplementary Figure S9.** Per-residue rmsf of SARS-CoV-2 RBD in complex with human ACE2 mutated at various residues in the 386AAQPFLL392 motif in silico. Mutagenesis was performed with the swapaa command in Chimera v. 1.13 and the structure was minimized with 2000 steps of steepest descent. Residues of interest in comparison to the wild-type SARS-CoV-2 RBD/hACE2 complex are labeled.

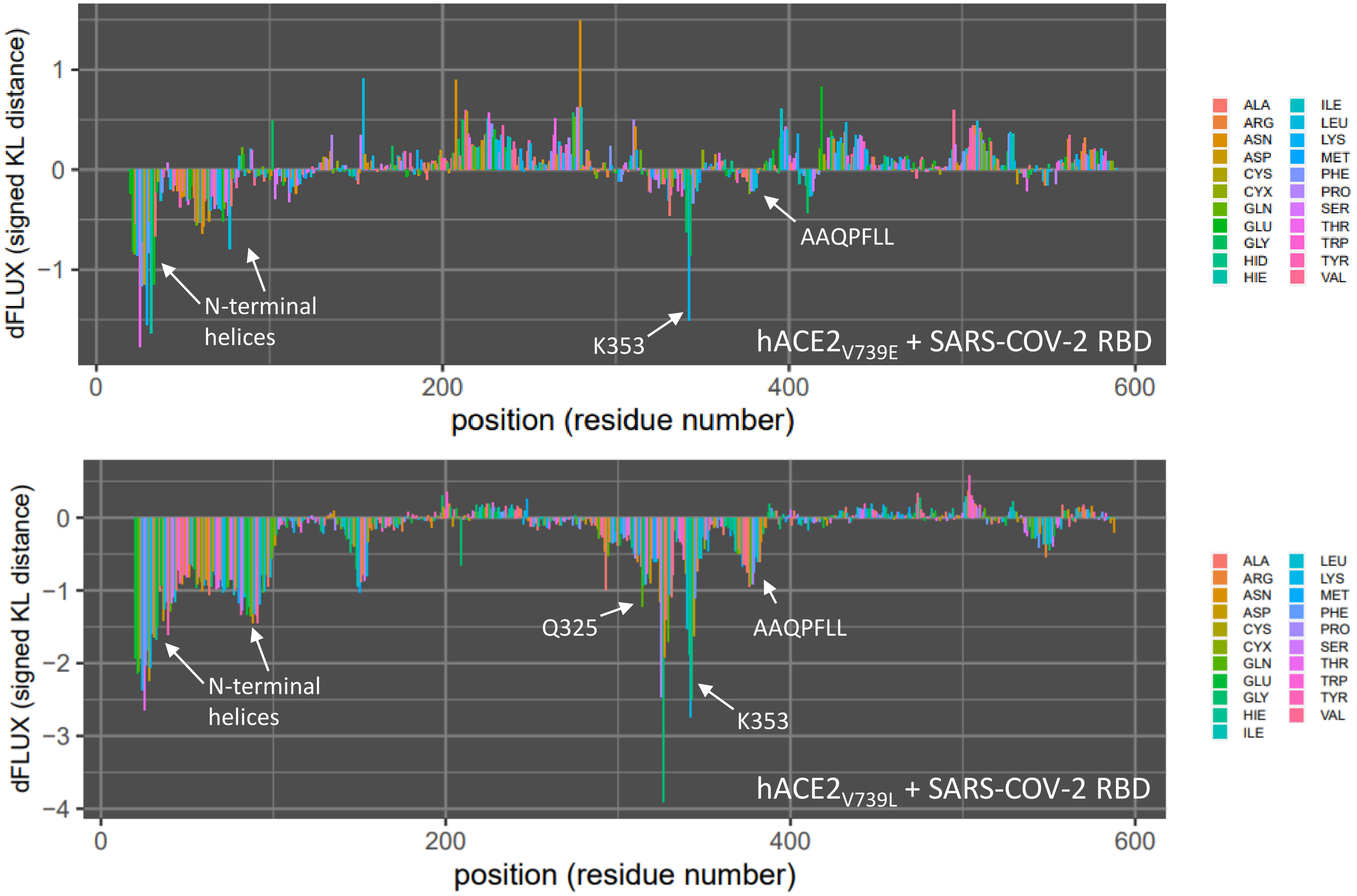


**Supplementary Figure S10.** Per-residue rmsf of SARS-CoV-2 RBD in complex with human ACE2 mutated at the V739 position to hydrophobic residue leucine and hydrophilic residue glutamate. Mutagenesis was performed with the swapaa command in Chimera v.1.13 and the structure was minimized with 2000 steps of steepest descent. Residues of interest in comparison to the wild-type SARS-CoV-2 RBD/hACE2 complex are labeled.

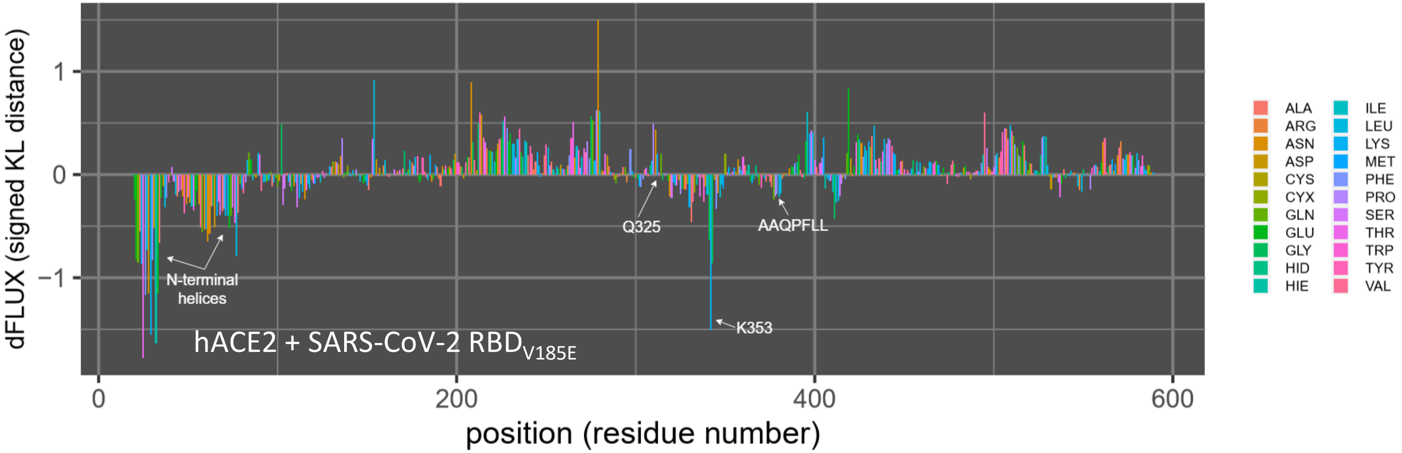


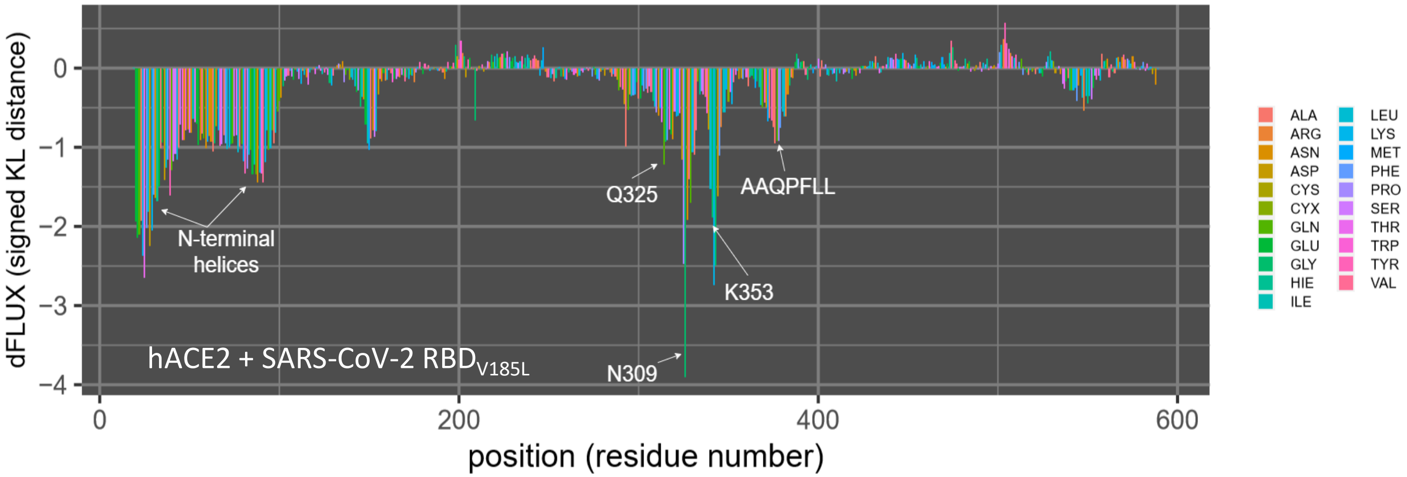


**Supplemental Figure S11.** Per-residue rmsf of ACE2 in complex with SARS-CoV-2 RBD mutated at the V185 position in silico. Mutagenesis was performed with the swapaa command in Chimera v. 1.13 and the structure was minimized with 2000 steps of steepest descent. Residues of interest in comparison to the wild-type SARS-CoV-2 RBD/ACE2 complex are labeled.

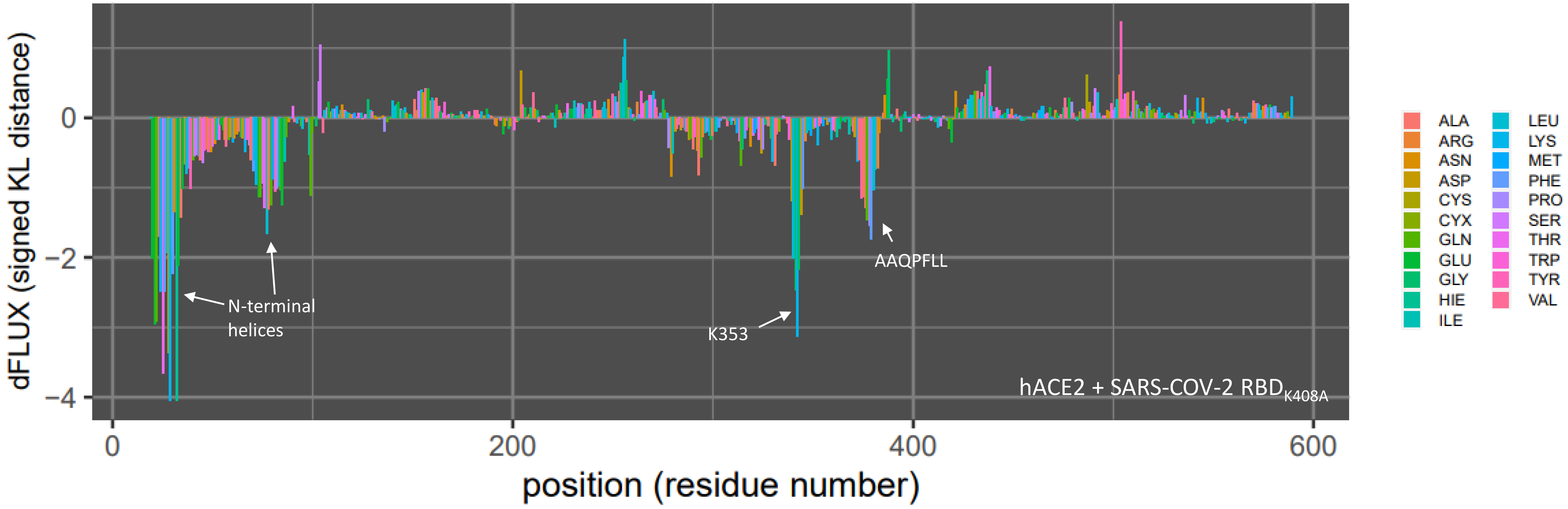


**Supplementary Figure S12.** Per-residue rmsf of SARS-CoV-2 RBD in complex with human ACE2 mutated at the K408 position to alanine. Mutagenesis was performed with the swapaa command in Chimera v. 1.13 and the structure was minimized with 2000 steps of steepest descent. Residues of interest in comparison to the wild-type SARS-CoV-2 RBD/hACE2 complex are labeled.
